# Loneliness as a modulator of intersubject neural alignment across social and nonsocial contexts

**DOI:** 10.64898/2026.11.03.763905

**Authors:** Bokyeong Jeong, M. Justin Kim

## Abstract

Loneliness is associated with cognitive biases that diverge from normative patterns of understanding and may be reflected in altered neural responses to the world. Here, we examined how loneliness modulates neural synchrony across individuals during the processing of social and nonsocial information. Using high-resolution movie-watching fMRI data (*n* = 178) from the Human Connectome Project, we applied intersubject correlation analyses to identify context-dependent neural signature of loneliness. Lonely individuals primarily showed more idiosyncratic responses in regions within the default mode network, visual association cortex, and amygdala during social contexts, and in reward-related circuitry during nonsocial contexts. This suggests less stereotyped processing of social information and reward in lonely individuals. In contrast, lonely individuals exhibited greater neural synchrony in the nucleus accumbens during social contexts and in somatomotor regions during nonsocial contexts. This potentially reflects shared experiences of social craving and engagement with a socially irrelevant physical environment. These effects were robust against demographic similarities, kinship, depression, and objective levels of social connection. Together, these findings identify loneliness as a modulator of intersubject neural alignment, which may contribute to shaping how individuals process and interpret different aspects of the world.

## Introduction

Defined as a distressing feeling due to unmet social needs, either quantitatively or qualitatively (Perlman & Peplau, 1981), loneliness is a transient experience in most cases but can be devastating to physical and psychological well-being when prolonged (Hawkley & Cacioppo, 2010; Qualter et al., 2015; Kim & Sul, 2023). Loneliness is associated with poorer sleep quality (Cacioppo et al., 2002), higher rates of cardiovascular diseases (Vasan et al., 2024), dementia (Lara et al., 2019), depression (Erzen & Çikrikci, 2018), suicidal ideation (McClelland et al., 2020), and greater mortality (Holt-Lunstad et al., 2015). Exacerbated by social phenomena that decrease face-to-face interaction, such as the COVID-19 pandemic, loneliness is becoming a growing concern around the world (Lam et al., 2021).

According to a model that explains the reinforcement of loneliness, cognitive bias serves as a core mechanism of loneliness and its detrimental consequences (Cacioppo et al., 2014). Lonely individuals suffer from cognitive biases during the stages of information processing, especially in social contexts (Cacioppo & Hawkley, 2009; Spithoven et al., 2017). For instance, lonely individuals tend to show heightened attention for social threats (Bangee et al., 2014; Qualter et al., 2013), expect rejection from others (Nowland et al., 2018; Watson & Nesdale, 2012), negatively attribute others’ intentions (Qualter et al., 2013), and evaluate themselves and others more negatively (Lodder et al., 2016).

Human neuroimaging studies have investigated whether lonely individuals’ biased information processing manifests as alterations in brain activity. For example, electroencephalogram (EEG) studies have shown that, relative to nonlonely individuals, lonely individuals differentiate socially threatening images more quickly than nonsocial threat images (Cacioppo et al., 2016), suggesting heightened implicit hypervigilance specifically toward social cues. A functional magnetic resonance imaging (fMRI) study found that lonely individuals show higher neural activity to people (vs. objects) in the visual cortex but lower activity to people (vs. objects) in the ventral striatum (Cacioppo et al., 2009), indicating a context-dependent differential pattern of visual and reward processing in lonely individuals. Other task-based fMRI studies employed images of humans that diverged in facial expression (Lindner et al., 2014; Wong et al., 2016) or distance of the relationship (Inagaki et al., 2016), as well as non-human images (Wong et al., 2016), to capture the context-dependent nature of cognitive biases in lonely individuals.

While highly controlled experimental paradigms used in these studies have been instrumental in demonstrating that lonely individuals exhibit cognitive biases predominant in the social contexts, this level of experimental control may come at the expense of ecological validity (Lam et al., 2021; Nastase et al., 2019). That is, in such experiments, behavioral and neural responses to social interactions are often assessed using static, decontextualized photographs. In contrast, everyday social experiences involve perceiving and interacting with both unfamiliar and familiar individuals within rich, dynamic, and contextually embedded environments. Considering loneliness is maintained and reinforced by complex interplay between external social experiences and individuals’ predisposed characteristics (Perlman & Peplau, 1981), incorporating more naturalistic stimuli may provide an important complement to highly controlled experimental paradigms.

Motivated by this, in this study, we take advantage of naturalistic paradigms and movie-watching fMRI data to explore whether context-dependent cognitive biases are reflected in the neural patterns of lonely individuals. Movie-watching fMRI data lends itself to intersubject correlation (ISC) analysis that indexes neural synchrony across people, which circumvents the need for a pre-defined response model by measuring similarity of neural responses between participants to dynamic, complex naturalistic stimuli (Hasson et al., 2004; Nastase et al., 2019). When participants exhibit high ISCs, we can infer several types of shared processing across participants during exposure to a naturalistic stimulus, such as similarities in attentional engagement (Song et al., 2021), memory encoding (Hasson et al., 2008), affective processing (Nummenmaa et al., 2012), and even high-level cognitive processing, including perspective taking (Lahnakoski et al., 2014) and interpretation (Yeshurun et al., 2017). By using this approach, we sought to gain insight into whether the context-sensitive cognitive processing features in lonely individuals can be captured during movie-watching that resembles real-world events.

An important prior study (Baek et al., 2023) has examined the ISC of lonely individuals with the assumption that loneliness, characterized by a lack of feeling understood (Routasalo et al., 2006), might be driven by the possibility that they perceive the world differently from others. Results showed that lonely individuals process the world in their own way (i.e., low ISCs), whereas nonlonely individuals are more alike to each other (i.e., high ISCs). However, considering the aforementioned context dependency of cognitive biases in lonely individuals, critical questions still remain. First, lonely individuals can see the world differently from others, but this idiosyncrasy may only occur within a social context, while lonely individuals see the world just the same as others in the nonsocial context. Moving beyond a purely holistic approach, incorporating the context-sensitive nature of cognitive biases in lonely individuals (Cacioppo & Hawkley, 2009; Cacioppo et al., 2009, 2016; Spithoven et al., 2017) may offer additional insight into the cognitive processing characteristics associated with loneliness. Second, cognitive biases of lonely individuals can be expressed as both idiosyncrasy and synchrony in brain responses. Since ISC is not identical to activation, any consistent increases and decreases of brain responses across participants can yield high ISCs (Nastase et al., 2019). While Baek and colleagues (2023) hypothesized that lower ISCs (i.e., idiosyncrasy) would be linked to higher loneliness, it is still possible that certain brain regions may show higher ISCs (i.e., synchrony) in lonely individuals. Together, we sought to extend previous findings that lonely individuals exhibit lower ISCs (Baek et al., 2023) by examining the context dependence and functional specificity of this effect, with the goal of developing a more nuanced understanding of cognitive biases associated with loneliness.

First, we hypothesized that the effect of loneliness on neural synchrony across participants would be context-dependent, reflecting cognitive biases in loneliness. In studies that adopted conventional paradigms, differential neural patterns for social and nonsocial images were compared to show the context-dependent nature of cognitive biases in lonely individuals (Cacioppo et al., 2009, 2016; Wong et al., 2016). Based on this framework, we sought to find evidence for distinct patterns of ISCs between social and nonsocial contexts in lonely individuals to explain the cognitive biases in loneliness.

Specifically, we expected that the observed idiosyncratic processing in lonely individuals in the previous finding (Baek et al., 2023) would be predominant in the social context. We expected a significant interaction between loneliness and context (i.e., social and nonsocial), suggesting that loneliness does not uniformly degrade neural synchrony across subjects in all aspects of the world, but rather modulates brain response similarity in a way that is sensitive to the social relevance of the stimuli.

According to prior findings, sharing the same psychological perspective leads to high ISCs (Lahnakoski et al., 2014), while a different interpretation of the same stimuli can induce low ISCs between participants (Yeshurun et al., 2017). Furthermore, lonely individuals show low ISCs in the default mode network (DMN), which is critical for understanding and mentalizing others’ intentions (Baek et al., 2023). Given these findings, we anticipated that lonely individuals would show idiosyncratic or less stereotypical patterns in the social context in higher-order brain areas associated with interpreting, attributing, and evaluating others’ behaviors, reflecting cognitive biases of loneliness prominent in social contexts (Cacioppo & Hawkley, 2009; Spithoven et al., 2017).

Second, although our primary *a priori* hypotheses focused on higher-order social cognition areas, we adopted a whole-brain, exploratory approach in order to investigate how loneliness might differentially modulate other functional systems across social and nonsocial contexts. We expected such idiosyncratic processing might be confined to social cognition areas in the social context, as any consistent and time-locked increase or decrease in neural activity while watching naturalistic stimuli can induce high ISCs (Finn et al., 2020; Nastase et al., 2019; Sonkusare et al., 2019). Thus, with respect to functional specificity, we explored whether neural synchronization could identify brain regions that reflect cognitive processing characteristics associated with loneliness. More broadly, the present study aimed to clarify which aspects of the world and which brain functions are particularly important for understanding the cognitive biases that characterize lonely individuals.

## Methods

### Participants

To investigate if and how lonely individuals’ context-sensitive cognitive biases are reflected in alterations in neural synchrony patterns, we leveraged the well-established dataset provided by the Human Connectome Project (HCP) S1200 Release (Van Essen et al., 2013). Participants were all healthy young adults aged 22-35 and engaged in a movie-watching paradigm during a high-resolution functional MRI scanning at 7T. Participants who missed at least one run out of four movie runs were dropped to ensure appropriate calculation of timeseries correlation between participants. Accordingly, the sample used in this work (*n* = 178) includes all available data for this paradigm.

### Movie-Watching fMRI Data

The HCP movie-watching fMRI data was acquired using a 7T Siemens Magnetom scanner with a 32-channel RF-receive head coil at the Center for Magnetic Resonance Research at University of Minnesota. All four movie runs were collected using a multiband gradient-echo-planar imaging (EPI) sequence with the following parameters: repetition time (TR) = 1000 ms, echo time (TE) = 22.2 ms, flip angle = 45 deg, field of view (FOV) = 208 × 208 mm, matrix = 130 × 130, spatial resolution = 1.6 mm^3^, number of slices = 85, multiband factor = 5, image acceleration factor (iPAT) = 2, partial Fourier (pF) sampling = 7/8, echo spacing = 0.64 ms, bandwidth (BW) = 1924 Hz/Px. The direction of phase encoding alternated between anterior-to-posterior (AP; MOVIE1, MOVIE4) and posterior-to-anterior (PA; MOVIE2, MOVIE3). Details regarding data acquisition and a minimal preprocessing pipeline for HCP data can be found elsewhere (Glasser et al., 2013; Van Essen et al., 2012).

Briefly, each run is 15 minutes long, consisting of four to five movie clips presented in a fixed order. Two runs (MOVIE1 and MOVIE3) are comprised of documentaries and independent films, whereas the other two runs (MOVIE2 and MOVIE4) include Hollywood movie clips. All fMRI data analyses were proceeded with FIX-denoised data, which completed standard preprocessing pipelines (including motion correction, distortion correction, high-pass temporal filtering, linear and nonlinear registration to MNI152 space, and removal of artifactual component timeseries plus regression of 24 confound timeseries using FSL’s MELODIC and FIX tool). Spatial smoothing with a 4.8mm full-width at half maximum Gaussian kernel was performed as recommended to improve temporal signal-to-noise ratio in ISC analysis (Pajula & Tohka, 2014) with high-resolution fMRI data (Triantafyllou et al., 2006). We extracted BOLD timeseries across the whole brain using 114 parcels: 100 cortical parcels from the Schaefer-Yeo cortical atlas (Schaefer et al., 2018) and 14 subcortical parcels from the Harvard-Oxford subcortical atlas.

### Behavioral Measures

In addition to the movie-watching fMRI scans, a wide range of behavioral measures were also collected from the HCP participants. In this work, our primary measure of interest was the loneliness survey from the NIH Toolbox on emotion (henceforth referred to as loneliness score), consistent with the existing studies on loneliness that used HCP data (Liégeois et al., 2019; Mwilambwe-Tshilobo et al., 2019; Yi et al., 2018). Using a median split of loneliness scores, we stratified our sample into a low-loneliness group (loneliness score ≤ 52.00, *n* = 91) and a high-loneliness group (loneliness score > 52.00, *n* = 89) (Fig. 1a). We then used linear mixed-effects model (explained below), formulated as a two-group contrast, to identify areas that are differentially synchronized according to loneliness levels.

**Fig. 1.**
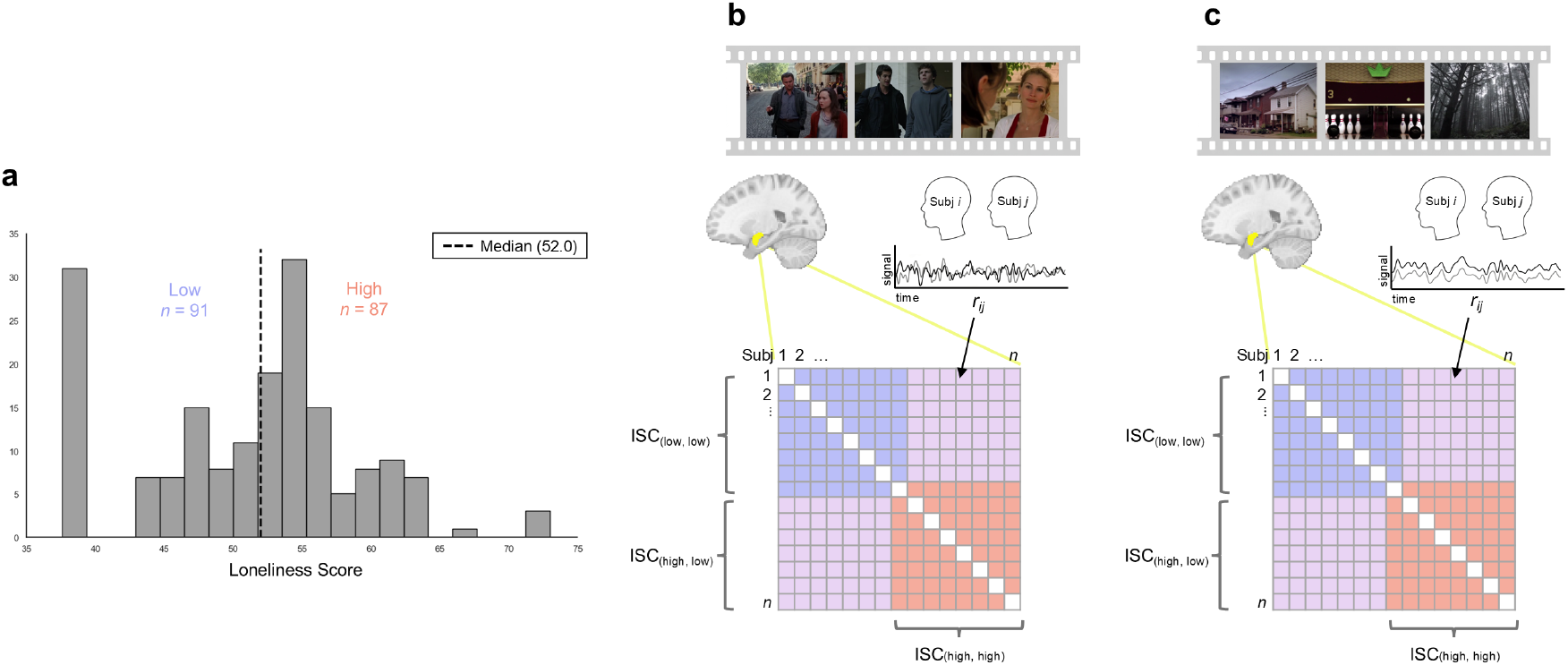
Distribution of loneliness scores and intersubject correlation (ISC) analysis. **a.** Distribution of scores on loneliness survey from the NIH Toolbox on emotion across a total of 178 participants. Median split is used to stratify participants into low (≤52, blue) and high (>52, red) loneliness. **b.** Schematic overview of ISC analysis in the social context. For each parcel, activation time course of a participant while viewing social scenes was regarded as a single timeseries, and correlated with another participant’s timeseries, yielding ISCs for each participant pair (*r_ij_*). This resulted in an ISC matrix for the social context for each parcel, in which the upper (or lower) off-diagonal reflects pairwise correlation coefficients of a unique number of participant pairs (*_178_C_2_* = 15,753). Then, these values were compared across loneliness groups using parcel-by-parcel linear mixed effects modelling with fully crossed random effects to account for the symmetric structure of the ISC matrix. **c.** Schematic overview of ISC analysis in the nonsocial context. For each parcel, activation time course of a participant while viewing nonsocial scenes was regarded as a single timeseries, and correlated with another participant’s timeseries, yielding ISCs for each participant pair (*r_ij_*). This resulted in an ISC matrix for the nonsocial context for each parcel, in which the upper (or lower) off-diagonal reflects pairwise correlation coefficients of a unique number of participant pairs (*_178_C_2_* = 15,753). Then, these values were compared across loneliness groups using parcel-by-parcel linear mixed effects modelling with fully crossed random effects to account for the symmetric structure of the ISC matrix.

Here, our primary focus was on loneliness, or subjective social isolation, which is conceptually distinct from objective social isolation (Hawkley et al., 2008). To account for the close and often inseparable relationship between subjective and objective social isolation, we additionally included the friendship survey from the NIH Toolbox on emotion – which assesses the perceived availability of companions for social interaction and affiliation – as a control variable in the analyses. Importantly, we sought to address that both behavioral and neural similarities can be affected by real-world intimacy of the participants (Hove & Risen, 2009; Liu et al., 2019; Miles et al., 2009; Parkinson et al., 2018). Since we expect both behavioral and neural similarity to be more greater among kin (Liu et al., 2019), we controlled for whether participants are siblings. In addition, a multitude of studies have reported on the significant link between loneliness and depression (Erzen & Çikrikci, 2018). In order to focus on cognitive processing features solely expressed by loneliness, we also included depression (Achenbach adult self-report DSM-oriented depressive problems scale) as a covariate. Lastly, we also included basic demographic variables (age, gender, and race), considering the diversity of HCP data (Van Essen et al., 2012) and consistency with prior studies on ISCs (Baek et al., 2023; Hyon et al., 2020). These steps ensured that the results were attributable to loneliness (see *Supplementary Materials* for details).

### Movie Stimuli and Event Segmentation

Given that movie stimuli comprise a complex mixture of multimodal sensory information, the four movies were annotated on a TR-by-TR basis using the video annotation research tool Anvil (http://www.anvil-software.org/) according to their visual and auditory features. With respect to visual features, scenes were annotated following a simple rule that was previously used in a study with a movie-watching paradigm (Wagner et al., 2016): ‘social’ defined as when two or more characters were interacting (visible or implied through action and/or conversation such as shot/reverse shot), ‘single’ defined as when a character was alone on the screen, ‘absent’ defined as when no characters were visible. To improve precision, the scenes in which two or more characters were present as mere juxtaposition were regarded as ‘multiple’. With respect to auditory features, we classified scenes in three different categories: ‘dialogue’ defined as when characters had a bidirectional conversation, ‘monologue’ defined as when one character talked unidirectionally, and ‘nobody talking’ defined as when any human verbalization was absent (see Table S1 for details of the stimuli).

To ensure the purity of the data, scenes with animals (e.g., dogs) or nonhuman creatures (e.g., Chewbacca from the movie *Star Wars*) were excluded, since there might be different views on whether they are social agents. As each movie consisted of four to five movie clips with 20 seconds of rest each in between the clips as well as at the start and end of the movies, we trimmed all the TRs that corresponded to rest. Importantly, movie clips were presented as if they were ‘blocks’ just as in a traditional GLM; if we were to include all TRs corresponding to movie clips, the BOLD response signal will be primarily driven by the gap between the rest and movie clips, distorting the ISCs (Nastase et al., 2019). As recommended by Natase et al. (2019), we also excluded an additional 10s at the onset of each epoch to mitigate this problem. Furthermore, within each functional run and each parcel, the timeseries was standardized and shifted backward in time by 5 TRs (i.e., 5s) to take into account the hemodynamic response delay (Hasson et al., 2004; Hasson et al., 2008).

Then, we concatenated all four functional runs into a single timeseries, and the combination of visual and auditory features was gathered to select TRs that represent social and nonsocial contexts. In this way, the scenes with a combination of ‘social’ visual feature with ‘dialogue’ auditory feature were sorted out, yielding the ‘social’ context that corresponded to 736 TRs, while the scenes with a combination of ‘absent’ visual feature with ‘monologue’ or ‘nobody talking’ auditory feature were selected as the ‘nonsocial’ context, which corresponded to 722 TRs (Fig. 2).

**Fig. 2.**
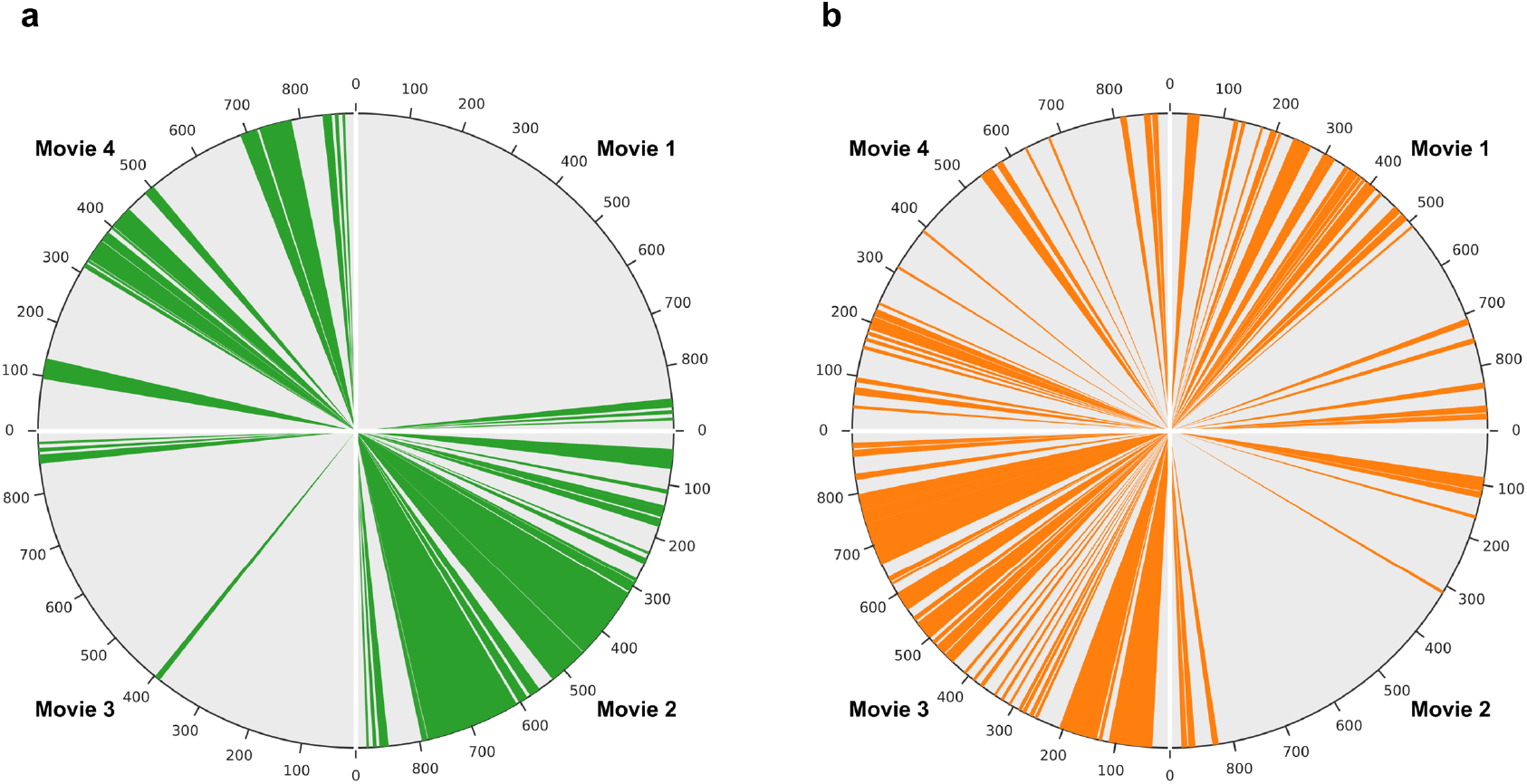
Movie stimuli and event segmentation. **a.** The timepoints selected to represent the social context. The four movie stimuli are shown clockwise, and scenes incorporating visual social interactions with auditory dialogue were selected, yielding 736 TRs. **b.** The timepoints selected to represent the nonsocial context. The four movie stimuli are shown clockwise, and scenes in which characters are visually absent, accompanied by either a monologue or no speech, were selected, yielding 722 TRs.

### Intersubject Correlation

We used ISC to analyze fMRI data, which is a model-free approach measuring the synchrony of the neural responses across participants in a given brain region. One participant’s response timeseries in one brain region is correlated with a second participant’s response timeseries at the same region. We consider a brain region with high ISC as having a shared or stereotyped functional role when processing the stimuli across subjects. For the ‘social’ context, we extracted average timeseries of voxels within each of the 114 parcel and calculated Pearson correlation coefficients for each pair of participants (Fig. 1b). As there were 178 participants, the total number of ISCs for a unique pair of participants was 15,753, which corresponds to either the lower or upper triangle of the off-diagonal of the matrix. ISC matrix for the ‘nonsocial’ context was generated in the same manner (Fig. 1c). Fisher’s *z*-transformation was applied to the ISC values prior to the main analyses.

### Linear Mixed Effects Modelling

First, we aimed to identify brain regions that showed significant differences in ISCs according to loneliness levels across the whole movie. In addition, we tested whether ISCs were significantly different between social and nonsocial contexts across all participants. To this end, we performed linear mixed effects modelling (LMM) as recommended by Chen et al. (2017). This approach effectively considers the symmetric nature of the ISC matrix by intentionally doubling the data to make fully crossed random effects. Due to the redundancy of the input data, we made manual adjustments from *2N – k* to *N – k* in degrees of freedom, where *N* is the number of unique pairs and *k* is the number of fixed effects in the model (Chen et al., 2017). We fit a linear mixed effects model with fully crossed random effects for each brain parcel using *LMER* and *LME4*, and performed planned contrasts presented below using *emmeans* in the R environment. We also fit a linear mixed effects model for each brain parcel, including control variables. All the statistical reports on the fMRI data were corrected for multiple comparisons using a false discovery rate at *p* < .05.

When investigating the effects of loneliness on ISCs, we were mainly interested in three contrasts. First, which brain regions show greater ISCs among pairs of high-loneliness subjects versus low-loneliness subjects, or vice versa (*ISC*_ℎℎ_ > *ISC_ll_*)? Second, which brain regions show greater ISCs among pairs of high-loneliness subjects versus mismatched pairs (i.e., high-low), or vice versa (*ISC*_ℎℎ_ > *ISC*_ℎ*l*_)? Lastly, which brain regions show greater ISCs among mismatched pairs versus pairs of low-loneliness subjects, or vice versa (*ISC*_ℎ*l*_ > *ISC_ll_*)? All three contrasts ask which brain regions show higher or lower ISCs in groups with more high-loneliness subjects than their counterparts during the processing of naturalistic stimuli.

In our main analysis, we investigated whether the ISC difference maps for each context were statistically different from each other. For instance, if ISC in the left amygdala were found to be significantly different between high-loneliness and low-loneliness pairs in the social context, while the difference was nonsignificant in the nonsocial context, this does not necessarily mean that the effect of loneliness on ISC in this region is significantly larger in the social context compared to the nonsocial context. In light of statistical guidelines cautioning against interpreting effects based solely on statistical significance in one condition and nonsignificance in another (Gelman & Stern, 2006), we directly tested whether brain regions whose intersubject correlations were modulated by loneliness in one context showed no modulation – or a qualitatively different pattern of modulation – by loneliness in the other context. Thus, we explored whether the interaction effect between loneliness and context was significant, performing LMMs with the aforementioned contrasts, both with and without the inclusion of the control variables.

Subsequently, we further probed which brain regions show significant differences in ISCs according to loneliness levels in social and nonsocial contexts, respectively. Using the ISC matrices for the social context (Fig. 1b) and the nonsocial context (Fig. 1c), we again performed LMMs using the same procedure described above, but this time separately for social and nonsocial contexts. This produced ISC difference maps with standardized beta coefficients for both social and nonsocial contexts, respectively.

## Results

### Descriptive Statistics

With respect to loneliness score, we observed a slightly left-skewed distribution with some variance (range = 37.60-73.00, mean = 50.94, S.D. = 8.17; median = 52.00, M.A.D. = 6.47; skewness = -0.09, see Fig. 1a for a histogram of the distribution). We opted for a median split to stratify participants into high- and low-loneliness groups rather than using the raw loneliness score as a continuous variable due to the unique challenge in pairwise ISC analysis (Fig. 1a). To make use of pairwise correlation values, other variables are required to be at the subject pair level as well, rather than the individual subject level. In addition, the influence of extreme values can be mitigated by a median split, avoiding an outsize effect on the result (Finn et al., 2018).

In our sample, loneliness score showed a significant negative relationship with the friendship survey from the NIH Toolbox on emotion (*ρ* = -.56, *p* < .001) and a significant positive relationship with Achenbach adult self-report DSM-oriented depressive problems scale (*ρ* = .35, *p* < .001). See Table S2 for a full report on descriptive statistics and bivariate associations.

### Main Effect of Loneliness

First, we investigated whether there was a main effect of loneliness on neural synchrony (i.e., ISC) when viewing the entire movie. We transformed individual-level loneliness score into pair-level (i.e., high-high, high-low, and low-low) and used planned contrasts with LMM to see if and how the ISCs in pairs with high-loneliness subjects show differences in each brain region. Among the three planned contrasts (*ISC*_ℎℎ_ > *ISC_ll_*; *ISC*_ℎℎ_ > *ISC*_ℎ*l*_; *ISC*_ℎ*l*_ > *ISC_ll_*), the left pallidum emerged as being more idiosyncratic (i.e., low ISCs) in the high-loneliness group than the group with mismatched pairs (β = -.04, *p* = .04). However, when controlling for the covariates of no interest, including demographic similarity, whether the pair is sibling, depression, and objective level of social connection, no single region remained significant in all three contrasts.

### Main Effect of Context

We also tested whether there was a main effect of context on neural synchrony across participants regardless of loneliness. Broad and bidirectional differences in brain synchrony were observed in 103 out of a total of 114 brain regions, in which ISCs significantly increased or decreased in either social or nonsocial contexts. See Table S3, S4 in the *Supplementary Materials* for a full list. Considering such distinct moderating effect of context on neural synchrony, we investigated the interaction between loneliness and context under the hypothesis that the effect of loneliness might depend on social vs. nonsocial contexts.

### Interaction Effect Between Loneliness and Context

Next, we aimed to identify brain regions whose neural synchrony was modulated by loneliness for one context but not modulated or differently modulated by loneliness level in the other context. Since comparing significance in one condition and nonsignificance in the other condition does not imply statistical significance *per se* (Gelman & Stern, 2006), we formally tested the interaction between loneliness and context (i.e., social and nonsocial context). We found several regions with significant interaction for the three planned contrasts (Fig. 3). As the beta value for the interaction effect quantifies the difference in the effects of loneliness in social and nonsocial contexts, it only signifies whether the modulating pattern of loneliness was significantly distinct between the two contexts in a given brain region. Thus, we further probed how the effect of loneliness on neural synchrony is manifested in each context after analyzing the interaction effect between loneliness and context.

**Fig. 3.**
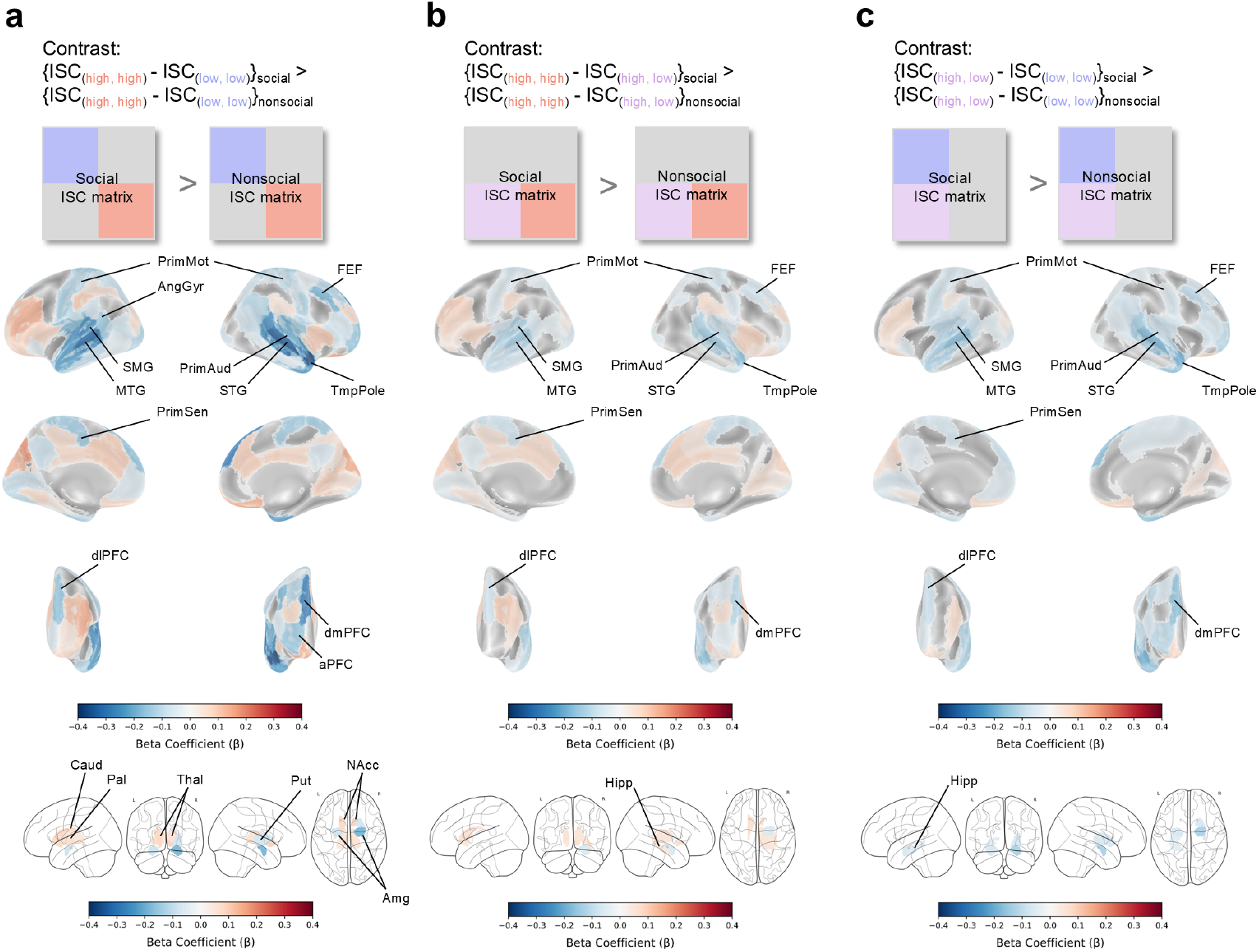
Interaction effect between loneliness and context. **a.** Schematized matrices indicate a contrast testing whether ISC difference between high-loneliness pairs and low-loneliness pairs (*ISC_hh_* > *ISC_ll_*) is significantly different in social compared to nonsocial context (top panel). Cortical results (middle panel) and subcortical results (bottom panel) are shown with a false-discovery-rate (FDR) corrected significance threshold of *p* < .05. **b.** Schematized matrices signify a contrast testing whether ISC difference between high-loneliness pairs and mismatched pairs (*ISC_hh_ > ISC_hl_*) is significantly different in social compared to nonsocial contexts (top panel). Cortical results (middle panel) and subcortical results (bottom panel) are shown with an FDR-corrected significance threshold of *p* < .05. **c.** Schematized matrices indicate a contrast testing whether ISC difference between mismatched pairs and low-loneliness pairs (*ISC_hl_ > ISC_ll_*) is significantly different in social compared to nonsocial contexts (top panel). Cortical results (middle panel) and subcortical results (bottom panel) are shown with an FDR-corrected significance threshold of *p* < .05. Each social and nonsocial ISC matrix is simplified to aid identification of the contrast. PrimMot = Primary motor cortex, AngGyr = Angular gyrus, SMG = Supramarginal gyrus, MTG = Medial temporal gyrus, FEF = Frontal eye fields, PrimAud = Primary auditory cortex, STG = Superior temporal gyrus, TmpPole = Temporal pole, PrimSen = Primary sensory cortex, dlPFC = dorsolateral prefrontal cortex, dmPFC = dorsomedial prefrontal cortex, aPFC = anterior prefrontal cortex, Caud = Caudate, Pal = Pallidum, Thal = Thalamus, Put = Putamen, NAcc = Accumbens, Amg = Amygdala, Hipp = Hippocampus.

In the first contrast (*ISC*_ℎℎ_ > *ISC_ll_*), where pairs of high-loneliness participants were compared with pairs of low-loneliness participants, we observed widespread interaction in 88 out of a total 114 parcels that spanned cortical and subcortical areas. In the cortical regions, the interaction effect was most notable in the right temporal pole, right superior temporal gyrus, left medial temporal gyrus, right dorsomedial prefrontal cortex, left supramarginal gyrus, right primary auditory cortex, right frontal eye fields, left primary sensory cortex, left dorsolateral prefrontal cortex, right anterior prefrontal cortex, left angular gyrus, and bilateral primary motor cortex. In the subcortical region, the interaction was significant in the bilateral amygdala, bilateral accumbens, left pallidum, left caudate, bilateral thalamus, and right putamen (for a complete list, see Fig. 3a and Table S5 in the *Supplementary Materials*).

Consistent with the first group contrast, the second and third contrasts (*ISC*_ℎℎ_ > *ISC*_ℎ*l*_, *ISC*_ℎ*l*_ > *ISC_ll_*) also revealed a broadly distributed interaction pattern across the brain, although the effects were somewhat attenuated in magnitude. These regions include the right temporal pole, right superior temporal gyrus, left medial temporal gyrus, right dorsomedial prefrontal cortex, left supramarginal gyrus, right primary auditory cortex, right frontal eye fields, left primary sensory cortex, left dorsolateral prefrontal cortex, and bilateral primary motor cortex (for a complete list, see Fig. 3b, 3c, and Table S6, S7 in the *Supplementary Materials*). In the subcortical regions, a similar pattern as in the first contrast (i.e., *ISC*_ℎℎ_ > *ISC_ll_*) was observed, with the right hippocampus (in the second contrast) and left hippocampus (in the third contrast) additionally emerging as having significant interaction between loneliness and context (for a complete list, see Fig. 3b, 3c, and Table S6, S7 in the Supplementary Materials).

Next, we also fit the model with covariates of no interest. We found a highly similar pattern of interaction effect between loneliness and context, even after controlling for pairwise similarity of demographic variables, kinship, depression, and self-reported levels of objective social connection (see Fig. S1 in the *Supplementary Materials*).

### Simple Main Effect of Loneliness in the Social Context

To further investigate context-specific effects of loneliness, we probed how groups show differences in neural synchrony for each brain region and in each context. We obtained ISC difference maps with standardized beta coefficients for social and nonsocial contexts, respectively. For each context, we first analyzed the data with the model in which ISCs in a particular brain region served as the dependent variable and loneliness groups as the independent variable (Fig. 4, 5, Table S8, S9), and then again analyzed with the model that included covariates of no interest (Fig. S2, S3). Note that not all regions with significant interaction emerged as being significantly modulated by loneliness in either or both contexts. Here, we focus on the brain regions where the interaction between loneliness and context was significant, as well as the simple main effect of loneliness at one context reached statistical significance when probed.

**Fig 4.**
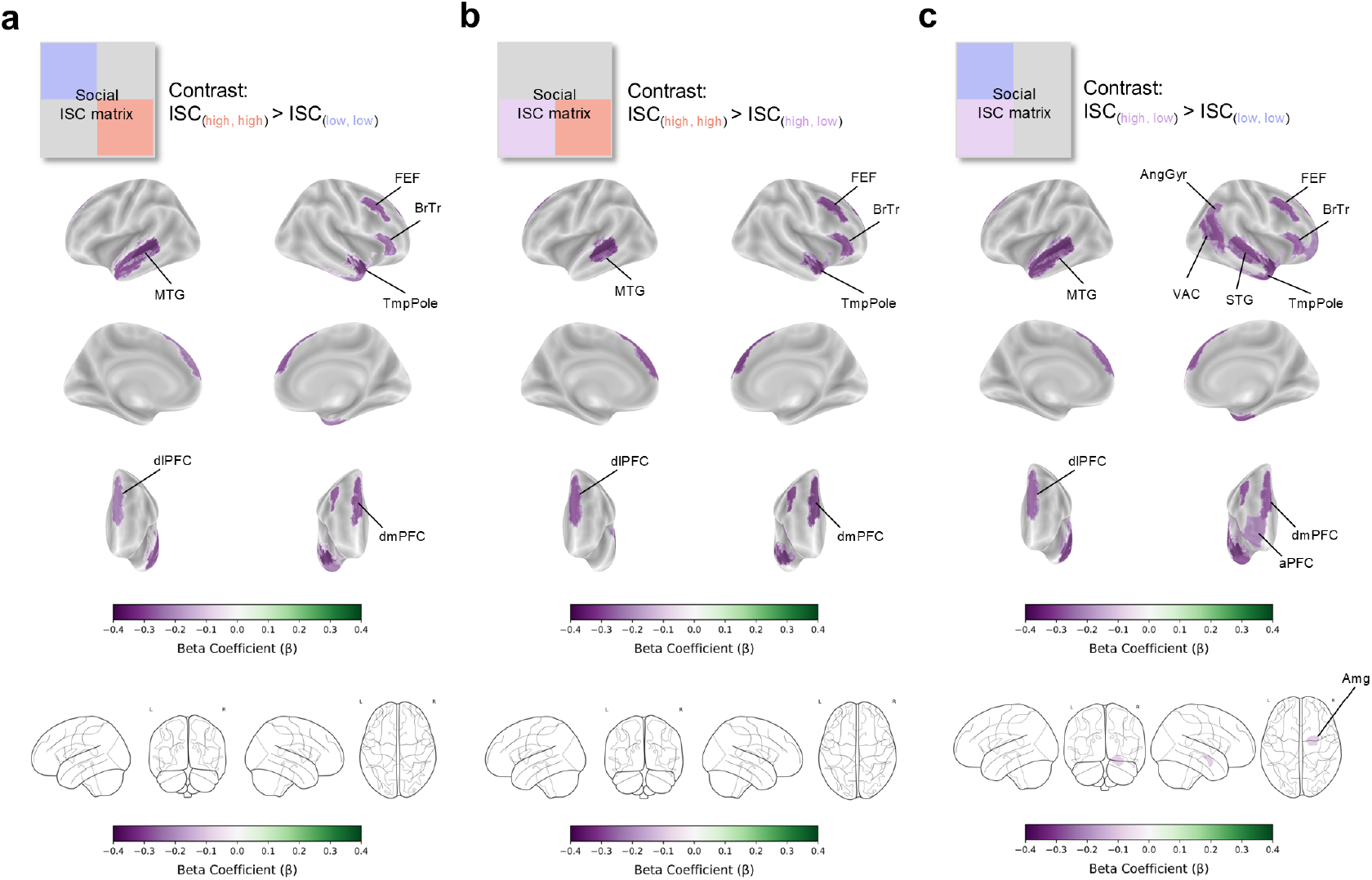
Simple main effect of loneliness in the social context. **a.** Schematized matrix indicates a contrast testing whether ISCs of high-loneliness pairs and low-loneliness pairs (*ISC_hh_* > *ISC_ll_*) are significantly different during viewing social context. (top panel). Cortical results (middle panel) and subcortical results (bottom panel) are shown with a false-discovery-rate (FDR) corrected significance threshold of *p* < .05. **b.** Schematized matrix signifies a contrast testing whether ISCs of high-loneliness pairs and mismatched pairs (*ISC_hh_ > ISC_hl_*) are significantly different while viewing social scenes (top panel). Cortical results (middle panel) and subcortical results (bottom panel) are shown with an FDR-corrected significance threshold of *p* < .05. **c.** Schematized matrix indicates a contrast testing whether ISCs of mismatched pairs and low-loneliness pairs (*ISC_hl_ > ISC_ll_*) are significantly different during viewing social context (top panel). Cortical results (middle panel) and subcortical results (bottom panel) are shown with an FDR-corrected significance threshold of *p* < .05. The social ISC matrix is simplified to aid identification of the contrast. MTG = Medial temporal gyrus, FEF = Frontal eye fields, BrTr = Broca’s triangle, TmpPole = Temporal pole, dlPFC = dorsolateral prefrontal cortex, dmPFC = dorsomedial prefrontal cortex, AngGyr = Angular gyrus, VAC = Visual association cortex, STG = Superior temporal gyrus, aPFC = anterior prefrontal cortex, Amg = Amygdala.

In the social context, groups including more high-loneliness participants than their counterparts showed lower ISCs in both cortical and subcortical regions (Fig. 4, Table S8). In the first and second contrasts (*ISC*_ℎℎ_ > *ISC_ll_*, *ISC*_ℎℎ_ > *ISC*_ℎ*l*_), where pairs of high-loneliness participants were compared with pairs of low-loneliness participants or pairs of mismatched participants, the former group showed more idiosyncrasy in brain responses in the right temporal pole, left medial temporal gyrus, right dorsomedial prefrontal cortex, left dorsolateral prefrontal cortex, right Broca’s triangle, and right frontal eye fields (Fig. 4a, 4b). In the third contrast (*ISC*_ℎ*l*_ > *ISC_ll_*), a similar pattern was observed, but several additional regions emerged as being more idiosyncratic in pairs of mismatched participants than in pairs of low-loneliness participants. These regions include the right superior temporal gyrus, right angular gyrus, right anterior prefrontal cortex, right visual association cortex, and right amygdala (Fig. 4c).

In addition, we fit a model including covariates of no interest to see if the effect of loneliness on ISCs persists after controlling for pairwise similarity of demographic variables, whether the pair is siblings, depression, and self-reported levels of objective social connection. Consistent with the model without covariates, groups including more high-loneliness participants than their counterparts primarily showed lower ISCs in regions within the DMN (Fig. S2). In the first contrast (*ISC*_ℎℎ_ > *ISC_ll_*), we found that brain responses in the social context were more idiosyncratic in high-loneliness group relative to low-loneliness group in the right temporal pole, left medial temporal gyrus, right dorsomedial prefrontal cortex, left dorsolateral prefrontal cortex, right Broca’s triangle, right frontal eye fields, right superior temporal gyrus, right angular gyrus, and right anterior prefrontal cortex (Fig. S2a). In the second contrast (*ISC*_ℎℎ_ > *ISC*_ℎ*l*_), brain responses of high-loneliness group were more idiosyncratic than the mismatched group in the right temporal pole, left medial temporal gyrus, right dorsomedial prefrontal cortex, left dorsolateral prefrontal cortex, right Broca’s triangle, and right frontal eye fields (Fig. S2b). In the third contrast (*ISC*_ℎ*l*_ > *ISC_ll_*), patterns were close to the first contrast (i.e., *ISC*_ℎℎ_ > *ISC_ll_*), but additional regions were found to be more idiosyncratic in mismatched group compared with low-loneliness group, including the left angular gyrus, right visual association cortex, and bilateral amygdala (Fig. S2c).

Additionally, the right accumbens emerged as being more synchronized in the high-loneliness group relative to the low-loneliness group (Fig. S2a). Importantly, these regions (except right Broca’s triangle), which emerged as being significant in probing specific patterns of the effect of loneliness on ISCs in the social context, did not show significant effect of loneliness in the nonsocial context. In other words, the observed interaction effect of these regions was primarily driven by the modulating pattern of loneliness in the social context.

### Simple Main Effect of Loneliness in the Nonsocial Context

In the nonsocial context, effect of loneliness on neural synchrony was bidirectional, either increasing ISCs in regions within somatomotor network (SMN) or decreasing ISCs in other brain regions (Fig. 5, Table S9). In the first contrast (*ISC*_ℎℎ_ > *ISC_ll_*), where pairs of high-loneliness participants were compared with pairs of low-loneliness participants, the former group showed more synchrony in brain responses in the left premotor and supplementary motor cortex, bilateral primary motor cortex and right insula, while more idiosyncrasy was observed in the right medial posterior prefrontal cortex, right orbitofrontal cortex, and left pallidum (Fig. 5a). Similarly, in the second contrast (*ISC*_ℎℎ_ > *ISC*_ℎ*l*_), brain responses of high-loneliness group were more synchronized in the bilateral primary motor cortex and right insula, while more idiosyncratic responses were found in the right medial posterior prefrontal cortex, left dorsal anterior cingulate cortex, right Broca’s triangle, left pallidum, left accumbens, and left caudate (Fig. 5b). In the third contrast (*ISC*_ℎ*l*_ > *ISC_ll_*), similar patterns were observed but the effect size was somewhat attenuated compared to the first and second contrasts, with the mismatched group being more synchronized in the left premotor and supplementary motor cortex, and right primary motor cortex, while also being more idiosyncratic in the left orbitofrontal cortex (Fig. 5c).

**Fig 5.**
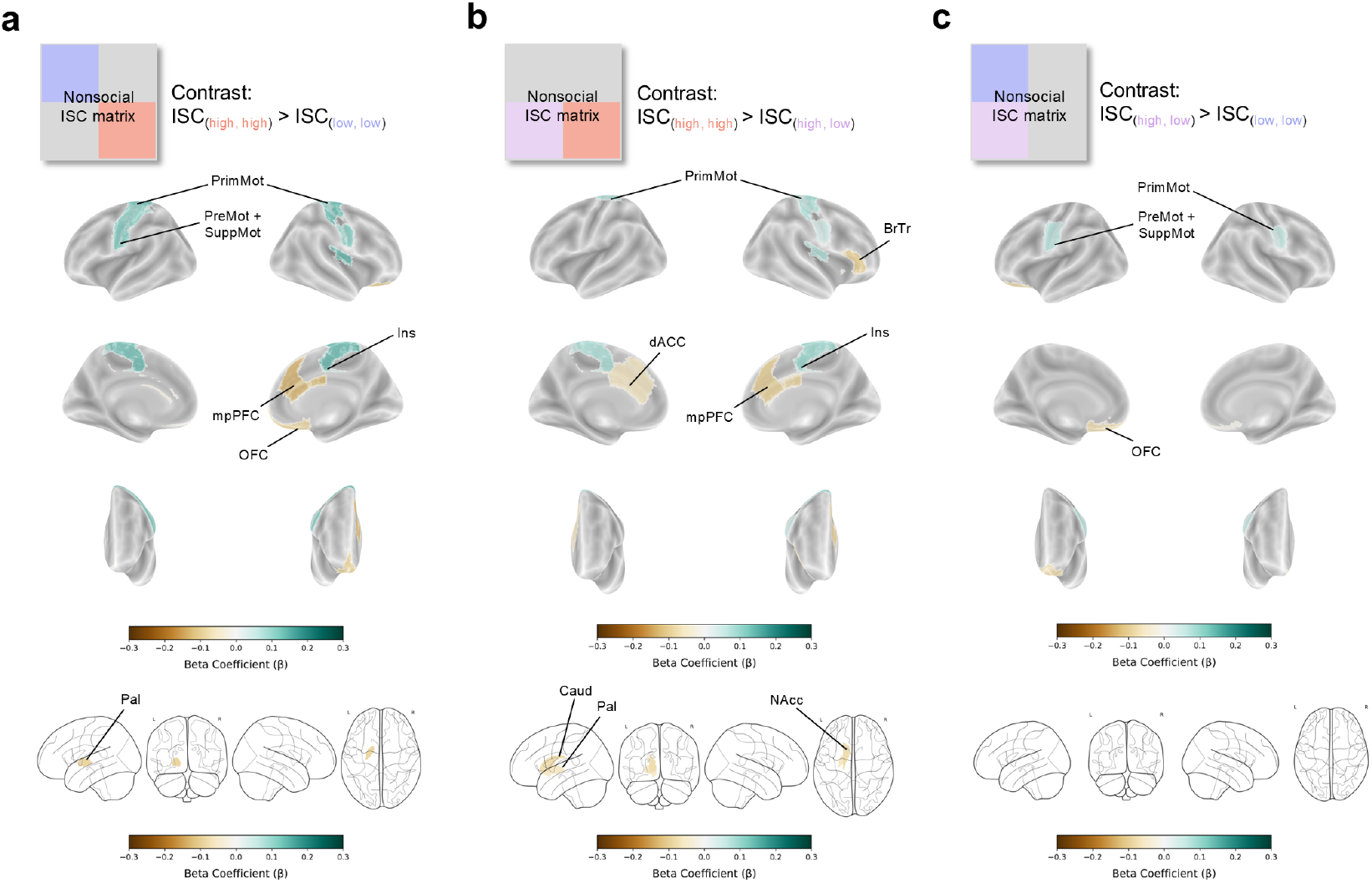
Simple main effect of loneliness in the nonsocial context. **a.** Schematized matrix indicates a contrast testing whether ISCs of high-loneliness pairs and low-loneliness pairs (*ISC_hh_* > *ISC_ll_*) are significantly different during viewing nonsocial context. (top panel). Cortical results (middle panel) and subcortical results (bottom panel) are shown with a false-discovery-rate (FDR) corrected significance threshold of *p* < .05. **b.** Schematized matrix signifies a contrast testing whether ISCs of high-loneliness pairs and mismatched pairs (*ISC_hh_ > ISC_hl_*) are significantly different while viewing nonsocial scenes (top panel). Cortical results (middle panel) and subcortical results (bottom panel) are shown with an FDR-corrected significance threshold of *p* < .05. **c.** Schematized matrix indicates a contrast testing whether ISCs of mismatched pairs and low-loneliness pairs (*ISC_hl_ > ISC_ll_*) are significantly different during viewing nonsocial context (top panel). Cortical results (middle panel) and subcortical results (bottom panel) are shown with an FDR-corrected significance threshold of *p* < .05. The nonsocial ISC matrix is simplified to aid identification of the contrast. PrimMot = Primary motor cortex, PreMot + SuppMot = Premotor and Supplementary motor cortex, Ins = Insula, mpPFC = medial posterior prefrontal cortex, OFC = Orbitofrontal cortex, Pal = Pallidum, BrTr = Broca’s triangle, dACC = dorsal anterior cingulate cortex, Caud = Caudate, NAcc = Accumbens.

When controlling for covariates of no interest, groups including more high-loneliness participants than their counterparts primarily showed higher ISCs in motor-related regions, whereas lower ISCs were observed in other areas, consistent with the model without control variables. (Fig. S3). In the first contrast (*ISC*_ℎℎ_ > *ISC_ll_*), we found that brain responses were more synchronized in the high-loneliness group relative to the low-loneliness group in the left premotor and supplementary motor cortex, and bilateral primary motor cortex, while more idiosyncrasy was observed in the right medial posterior prefrontal cortex and left pallidum. (Fig. S3a). In the second contrast (*ISC*_ℎℎ_ > *ISC*_ℎ*l*_), high-loneliness group showed more synchrony than mismatched group in the bilateral primary motor cortex and right insula, whereas more idiosyncratic brain responses were observed in the right medial posterior prefrontal cortex, left dorsal anterior cingulate cortex, right Broca’s triangle, left pallidum, and left caudate (Fig. S3b). A similar pattern was observed in the third contrast (*ISC*_ℎ*l*_ > *ISC_ll_*), with the left premotor and supplementary motor cortex, and bilateral primary motor cortex being more synchronized, while the left orbitofrontal cortex being more idiosyncratic in the mismatched group relative to low-loneliness group (Fig. S3c).

Importantly, these regions (except right Broca’s triangle), which were found to be significant in probing specific patterns of the effect of loneliness on ISCs in the nonsocial context, did not emerge as having significant effect of the loneliness in the social context. In other words, the observed interaction effect of these regions was primarily driven by the modulating pattern of loneliness in the nonsocial context.

## Discussion

Our results demonstrate that lonely individuals do not process the world in a uniformly idiosyncratic manner; rather, their patterns of processing vary depending on the contextual environment in which they are situated. In our study, loneliness was associated with dissociable patterns of neural synchrony across social and nonsocial contexts. Notably, there was no single region that showed a significant effect of loneliness on brain response similarity across the whole movie, regardless of the context. In contrast, the distinction between social and nonsocial contexts emerged as a key factor modulating intersubject brain synchrony. These findings offer a foundation for investigating the possibility that the effect of loneliness is manifested in a context-sensitive manner.

In the social context, brain regions that lonely individuals showed dissimilar responses to their peers include the right temporal pole, left medial temporal gyrus, right dorsomedial prefrontal cortex, left dorsolateral prefrontal cortex, right Broca’s triangle, right frontal eye fields, right superior temporal gyrus, right angular gyrus, and right anterior prefrontal cortex. Interestingly, these regions primarily belong to the DMN, based on the Yeo-7 networks parcellation (Yeo et al., 2011). DMN is known as a sense-making network that actively and dynamically integrates external social information and preexisting internal self-related knowledge (Yeshurun et al., 2021). Prior studies consistently reported that increased brain response similarity in regions within the DMN reflects various types of shared understanding (Ohad et al., 2025), including similar interpretation (Finn et al., 2018; Nguyen et al., 2019; Saalasti et al., 2019; Yeshurun et al., 2017), emotional responses (Smirnov et al., 2019), humor perception (Jääskeläinen et al., 2016), socio-sexual desire (Chen et al., 2020), and even moral decision-making (Tei et al., 2019; van Baar et al., 2019). In light of these findings, the decreased alignment in neural responses across lonely individuals in regions within the DMN during the social context indicates a divergence in how they process rich, complex social cues.

In addition, our results dovetail with recent research on loneliness and ISCs (Baek et al., 2023), which observed lonely individuals having idiosyncratic functional brain responses with regions particularly in the DMN. Furthermore, prior research using UK Biobank data also reported that the DMN has the most consistent association with loneliness across three brain imaging modalities, suggesting the possibility of up-regulation of the DMN supporting mentalizing, imagination, and reminiscence to complement the social void in lonely individuals (Spreng et al., 2020). Notably, we found that this pattern remained significant even after controlling for demographic similarities, kinship, depression, and self-reported objective social connection. Thus, our results suggest that idiosyncratic or less stereotypical brain responses observed in the DMN, especially in the social context, may reflect divergent social information processing, serving as a neural marker of loneliness.

Furthermore, lonely individuals also showed idiosyncratic neural responses in the visual association cortex (VAC) and amygdala when viewing social scenes. The VAC plays a key role in modulating top-down visual attention engagement (Gazzaley et al., 2007), while the amygdala is involved in threat detection/orientation and emotional processing (LeDoux, 2007). The diminished neural response similarities in these regions with high-loneliness participants demonstrate that top-down attentional modulation and processing of emotionally charged social cues are less stereotypical in lonely individuals. This may reflect lonely individuals’ tendency to suffer from implicit hypervigilance or heightened attention in the social environment (Bangee et al., 2014; Cacioppo et al., 2016; Qualter et al., 2013).

Additionally, we found the right accumbens (NAcc) to have increased alignment in neural responses for lonely individuals during social context when controlling for covariates. According to an evolutionary perspective, loneliness has an adaptive value in that it promotes behavior to increase the likelihood of survival (Cacioppo et al., 2014). Just as hunger motivates us to eat, loneliness motivates us to seek social relationships. Based on this idea, a few studies reported associations between loneliness and NAcc that supports the perspective on loneliness as a state of social hunger. One study that questioned whether physical hunger and social hunger are processed similarly in the brain showed NAcc as the critical region that differentiates social hunger from physical hunger, suggesting a link between NAcc and craving for social interaction in lonely individuals (Tomova et al., 2020). Another study showed that lonely individuals had greater NAcc activity in response to images of close others relative to strangers, which was interpreted as yearning for social connections in lonely individuals (Inagaki et al., 2016). Taking these findings into account, one possible explanation for the observed neural synchrony in NAcc across lonely individuals is that it may reflect craving for social interactions, characterized in loneliness as a state of social hunger.

Crucially, this altered neural response similarity observed in the social context for high-loneliness participants was indeed context-dependent, as all regions also exhibited significant interaction between loneliness and context. Since they did not show significant group difference in the nonsocial context (except Broca’s triangle), it is evident that the effect of loneliness on neural synchrony in regions within the DMN, VAC, amygdala, and NAcc is almost exclusively driven by the social context. In other words, both idiosyncratic processing within DMN, VAC, and amygdala, and synchronized responses in NAcc across lonely individuals were uniquely triggered by the social context.

On the other hand, in the nonsocial context, functionally dissociable patterns of neural synchrony and idiosyncrasy were found according to loneliness levels. Interestingly, we found that lonely individuals showed similar neural responses to their peers in regions that are categorized to be a part of the SMN. This raises the possibility that lonely individuals tend to “tick collectively” to stimuli that deviate from social relevance (e.g., objects, landscapes). According to prior studies, it has been reported that simply viewing artifacts can activate motor-related imagery, thereby activating areas involved in movements, including the premotor cortex and supplementary motor cortex (Chao & Martin, 2000; Mizuguchi et al., 2013). Although these studies used a specific category of objects (i.e., man-made tools), the stimuli in this work incorporate considerably frequent appearances of artifacts such as a door handle or bowling balls. Those commonly associated with manual movements might have triggered motor-related imagery (Chao & Martin, 2000), thereby inducing increased alignment in neural responses observed in lonely individuals. Another possibility comes from the perspective of multi-sensory processing in motor-related areas. Prior studies reported that auditory, visual, and somatosensory cross-modality integration occurs in ventral premotor cortex (Driver & Noesselt, 2008), while supplementary motor areas are involved in auditory processing and auditory imagery (Lima et al., 2016). Considering the multi-modal complexity of the movie stimuli, the observed synchrony in regions within the SMN in lonely individuals might suggest increased cross-modal processing similarities when social cues are absent in the environment. Taken together, our findings show that lonely individuals do not always show idiosyncratic processing, but rather display shared, more stereotypical responses in the context lacking social relevance, perhaps reflecting time-locked, collective, and consistent motor-related imagery of objects and similar multi-sensory integration.

At the same time, lonely individuals showed idiosyncrasy in regions that constitute part of the brain’s reward system while watching nonsocial scenes. These regions include medial posterior prefrontal cortex (mpPFC), involved in top-down inhibitory control (Hester et al., 2009) and reward prediction to facilitate adaptation to the environment (Danielmeier et al., 2011; Hester et al., 2008; Ridderinkhof et al., 2004), orbitofrontal cortex (OFC), which is associated with updating reward values (Rudebeck & Rich, 2018), dorsal anterior cingulate cortex (dACC), processing reward-based decisions (Chau et al., 2018), pallidum and caudate that encode hedonic and motivational incentives (Castro et al., 2015; Haruno et al., 2004). These regions, belonging to the cortical-basal ganglia circuit as a core reward system in the brain (Haber & Knutson, 2010), showed idiosyncrasy in lonely individuals when viewing nonsocial scenes. This suggests loneliness might alter reward processing at the neural level, especially in the nonsocial context. This was evident in a prior study that demonstrated lonely individuals showed stronger activation to pleasant nonsocial images relative to pleasant social images, while participants with low loneliness levels showed the opposite pattern (Cacioppo et al., 2009). One possible explanation for this pattern was that lonely individuals may be left to find relative comfort in nonsocial than social cues (Cacioppo et al., 2009). In addition, studies on anthropomorphism demonstrated that loneliness influences how people understand a variety of nonsocial stimuli (Epley et al., 2007; Epley et al., 2008a, 2008b). Given these findings, the observed neural idiosyncrasy in reward circuitry in lonely individuals may indicate that lonely individuals do not find the same aspects of the nonsocial stimuli rewarding as their peers, reflecting divergent preferences and interpretations, such as anthropomorphism for nonsocial stimuli in lonely individuals.

In addition, we found idiosyncratic brain responses in the right Broca’s triangle in the nonsocial context as well as in the social context. The right-hemispheric homologue of Broca’s triangle plays supportive role in language production and comprehension (Naeser et al., 2011; Van Ettinger-Veenstra et al., 2010), specializing in affective prosody perception (Ross & Monot, 2008; Witteman et al., 2011). As both social and nonsocial scenes required linguistic processing, our result may suggest divergent processing of emotional intonation of utterances in lonely individuals regardless of the contexts. This finding supports prior findings that loneliness not only modulates recognition accuracy of emotional component in vocal expression (Morningstar et al., 2020; Shin & Kim, 2019) but also characterized by dissimilar language use, deviating from cultural consensus (Broom et al., 2024). Notably, the attenuating effect of loneliness on the neural synchrony was significantly more pronounced in the social context, as evidenced by the significant interaction between loneliness and context in this region. This may be partly attributed to more linguistic features being present (e.g., dialogue) in the social context.

Importantly, this altered neural synchrony found in the nonsocial context for high-loneliness participants was context-dependent, as all these regions also exhibited significant interaction between loneliness and context. Since they did not show a significant group difference in the social context (except Broca’s triangle), it is evident that the effect of loneliness on brain response synchrony in regions within the SMN and reward circuitry is almost exclusively driven by the nonsocial context. This reveals both synchronized processing within SMN and idiosyncratic responses in reward circuitry across lonely individuals were selectively triggered by the nonsocial context.

Our results have implications for explaining which aspects of the world and which brain functions are particularly critical in explaining lonely individuals’ cognitive processing. Loneliness modulates neural synchrony across participants in a context-dependent, functionally dissociable manner. Lonely individuals shared an instinctive craving for social interaction, but their attentional, emotional, and higher-order sense-making for the social cues were divergent in the social context. On the other hand, when facing socially detached cues, they showed idiosyncratic reward processing, while engagement with the physical features was more stereotypical. Therefore, we conclude that loneliness selectively reshapes neural responses according to contexts.

Moving beyond a holistic perspective, our findings highlight the importance of considering contexts to fully understand the cognitive processing features in lonely individuals. Future work can further expand on how lonely individuals respond to positive, negative, or neutral social and nonsocial naturalistic stimuli by taking into account their emotional valence. Moreover, to ensure objectivity, we annotated the visual and auditory features of the scenes exactly as they were, based on strict criteria agreed upon by the researchers, and then combined these to form social and nonsocial contexts. However, as the stimuli were edited, we only assume that they resemble real-world social or nonsocial contexts – we cannot guarantee how closely they resemble real-world events. Future work could complement this by exploring the similarities in brain responses of lonely individuals while participants are communicating or exposed to nonsocial stimuli through hyperscanning to capture brain-to-brain coupling during real-world events (Montague et al., 2002).

Furthermore, we acknowledge that it is difficult to distinguish whether the observed neural alterations in lonely individuals are the cause or consequence of loneliness. Nevertheless, we suggest that these unique cognitive processing features can be part of the self-reinforcing loop, being a core mechanism of maintaining and deteriorating loneliness (Cacioppo & Hawkley, 2009; Hawkley & Cacioppo, 2010). Lastly, it is important to note that intersubject correlation is conceptually different from an increase or decrease in brain activity. Future research could focus on how the unique cognitive processing patterns of lonely individuals manifest specific directions of activity to certain characteristics of naturalistic stimuli.

In summary, our findings suggest that loneliness may shape neural synchrony in a context-dependent and functionally dissociable manner. Lonely individuals show idiosyncratic responses in regions within DMN, VAC, and amygdala in the social context, reward circuitry in the nonsocial context, and Broca’s triangle in both contexts. This supports divergent and less stereotypical social information processing, nonsocial reward processing, and linguistic processing in lonely individuals. Moreover, atypical synchrony was found in the NAcc in the social context, and regions within the SMN in the nonsocial context for lonely individuals, reflecting the shared experience of social hunger and of a socially irrelevant physical world. Importantly, these effects remained significant even after controlling for demographic similarities, kinship, depression, and objective social connection. These findings lend support to the view that loneliness is a critical factor for modulating how we process the world, in ways that are both context-dependent and functionally dissociable.

## Declarations

## Acknowledgements

Data were provided by the Human Connectome Project, WU-Minn Consortium (Principal Investigators: David Van Essen and Kamil Ugurbil; 1U54MH091657) funded by the 16 NIH Institutes and Centers that support the NIH Blueprint for Neuroscience Research; and by the McDonnell Center for Systems Neuroscience at Washington University.

## Ethics Approval

Not applicable.

## Consent to Participate

Not applicable.

## Consent for Publication

Not applicable.

## Availability of Data and Materials

The dataset is publicly available at https://balsa.wustl.edu/ (HCP-Young Adult).

## Code Availability

The full code for the analyses will be shared at https://github.com/humanaffectneurolab upon publication.

