## Supplementary Materials for "Loneliness as a modulator of intersubject neural alignment across social and nonsocial contexts"

### Supplementary Methods

#### Computation of the Control Variables

We used linear mixed effects modelling to investigate the relationship between loneliness and intersubject correlation and added covariates of no interest to control for the similarities in demographic variables (age, gender, and race), kinship, depression, and objective level of social connection. To deal with a unique challenge in pairwise ISC analysis, we computed these individual-level variables into pair-level similarity measures.

With respect to age, we calculated the absolute value of the difference between the ages of the two individuals in each pair. Then, these values were transformed into a similarity measure, so that larger numbers indicate greater similarities (i.e., $age_{similarity} = 1 - \frac{age_{difference}}{\max(age_{difference})}$). To control for the self-reported gender, race, and kinship, we generated indicator variables for each so that 0 signifies a different gender, race, and mother ID, and 1 signifies the same gender, race, and mother ID. To control for self-reported levels of depression and objective levels of social connection, we took the value of the more distressed participant as the value of each pair. Rather than simply taking the average of the values for each pair, this approach allows not only to avoid information loss in pairs, but also to test the hypothesis in a consistent manner, testing the influence of the inclusion of one distressed participant (i.e., high-loneliness, high-depression, and low-objective social connection) in a pair on ISCs. This choice is consistent with the recent study that related neural similarity with behavioral measures (Baek et al., 2023). We used Achenbach adult self-report DSM-oriented depressive problems scale, taking the maximum value in each pair to represent the value for the pair. Similarly, we used friendship survey from the NIH Toolbox on emotion as a measure of objective levels of social connection, taking the minimum value in the pair as the value of each pair.

**Table S1.** Details of the stimuli.

| Movie No. | Clip No. | Title | Duration (s) | Contents |
|---|---|---|---|---|
| 1 | 1 | *Two Men* | 244 | A portrayal of two men running across a field under the scorching sun, and a man contemplating their intentions. Based on a short story, 'The Men Running Fast' by Franz Kafka. |
| | 2 | *Welcome to Bridgeville* | 222 | A description of people living in the town called Bridgeville and the history of the town. |
| | 3 | *Pockets* | 188 | People of various races and genders encountered on the street take turns explaining the items in their pockets and what they mean to them. |
| | 4 | *Inside the Human Body* | 64 | People who have overcome physical limitations appear one after another. |
| | 5 | *Vimeo Repeat* | 83 | Various natural landscapes, artificial objects, and people appear briefly. |
| 2 | 1 | *Inception* | 227 | Dom Cobb and Ariadne walk together through the depths of Cobb's subconscious. As they converse, the environment around them shifts and transforms with their imagination. |
| | 2 | *Social Network* | 259 | Focuses on Mark Zuckerberg's college life, specifically the controversy surrounding the website he built on campus, his appearance on the university's disciplinary board, and his relationship with peers. |
| | 3 | *Ocean's Eleven* | 249 | Danny Ocean gathers his potential accomplices in a house to unveil and discuss his heist plan. |
| | 4 | *Vimeo Repeat* | 83 | – |
| 3 | 1 | *Off the Shelf* | 181 | The story follows the journey of a red potted flower as it passes through the hands of various people and travels across different places. |
| | 2 | *1212* | 185 | A man and a woman find themselves in the same hotel but in different rooms. They notice the presence of each other by the faint sounds. |
| | 3 | *Mrs. Meyers Clean Day* | 204 | A woman talks about the reason she started the urban training farm and community edible garden, and its positive effects. |
| | 4 | *Northwest Passage* | 143 | A bleak rural landscape with a backdrop of dark music. |
| | 5 | *Vimeo Repeat* | 83 | – |
| 4 | 1 | *Home Alone* | 232 | Kevin wakes up in the morning and finds out that he is alone in his home. |
| | 2 | *Erin Brockovich* | 230 | Erin Brockovich takes her children to visit the victim and appeals for her to stand trial. |
| | 3 | *Star Wars: Episode V - The Empire Strikes Back* | 255 | Han Solo returns to the base and informs the general that he will soon be leaving. He engages in an argument with Princess Leia, questioning the sincerity of her love for him. |
| | 4 | *Vimeo Repeat* | 83 | – |

*Note.* 'Vimeo Repeat' is an edited clip including 'J.L. Powell 23 Degrees South' and 'LXIV'.

**Table S2.** Descriptive statistics and bivariate associations.

| | | Total Participants ($n$ = 178) | Low Loneliness ($n$ = 91) | High Loneliness ($n$ = 89) | Group Difference | Correlation |
|---|---|---|---|---|---|---|
| Age | | 29.40 (3.32) | 29.45 (3.36) | 29.34 (3.29) | $t$ = 0.21 ($p$ = .83) | $\rho$ = -.02 ($p$ = .82) |
| Gender | Female | 108 (60.67%) | 50 (54.95%) | 58 (66.67%) | $\chi^2$= 2.56 ($p$ = .11) | – |
| | Male | 70 (39.33%) | 41 (45.05%) | 29 (33.33%) | | |
| Race | Asian/Nat. Hawaiian/Othr Pacific Is. | 7 (3.93%) | 4 (4.40%) | 3 (3.45%) | $\chi^2$= 2.13 ($p$ = .55) | – |
| | Black or African Am. | 13 (7.30%) | 7 (7.69%) | 6 (6.90%) | | |
| | White | 156 (87.64%) | 78 (85.71%) | 0 (0.00%) | | |
| | Unknown or Not Reported | 2 (1.12%) | 2 (2.20%) | 78 (89.66%) | | |
| Friendship | | 50.05 (8.54) | 53.89 (7.49) | 46.03 (7.72) | $t$ = 6.90 ($p$ < .001) | $\rho$ = -.56 ($p$ < .001) |
| Depression | | 52.85 (4.52) | 51.69 (2.87) | 54.07 (5.52) | $t$ = -3.58 ($p$ < .001) | $\rho$ = .35 ($p$ < .001) |

*Note.* Each row is covariates of no interest – age, gender, race, friendship survey from the NIH Toolbox on emotion (henceforth friendship), and Achenbach adult self-report DSM-oriented depressive problems scale (henceforth depression). The former three columns indicate mean and standard deviation (with respect to age, friendship, and depression) or frequency and proportion (with respect to gender and race) for total participants, low- and high-loneliness participants. The fourth column shows a statistical comparison of the group differences according to the median split of the loneliness survey from the NIH Toolbox on emotion (Student's $t$-test or chi-square test). The last column indicates Spearman's correlation between loneliness score and each continuous variable. Asian/Nat. Hawaiian/Othr Pacific Is. = Asian/Native Hawaiian/Other Pacific Islander, Black or African Am. = Black or African American, Friendship = Friendship survey from the NIH Toolbox on emotion, Depression = Achenbach adult self-report DSM-oriented depressive problems scale.

**Table S3.** Brain regions showing significant main effect of context.

| Parcel No. | Hemisphere | Network | Cortical Results<br>Component | $ISC_{social}$ | $ISC_{nonsocial}$ | β | *SE* | *p* |
|---|---|---|---|---|---|---|---|---|
| 1 | Left | Vis | Visual | 0.229 | 0.173 | 0.597 | 0.005 | 0.000 |
| 2 | Left | Vis | Visual | 0.300 | 0.264 | 0.303 | 0.005 | 0.000 |
| 3 | Left | Vis | Visual | 0.311 | 0.229 | 0.669 | 0.005 | 0.000 |
| 4 | Left | Vis | Visual | 0.219 | 0.269 | -0.423 | 0.005 | 0.000 |
| 5 | Left | Vis | Visual | 0.305 | 0.325 | -0.149 | 0.005 | 0.000 |
| 6 | Left | Vis | Visual | 0.241 | 0.182 | 0.530 | 0.005 | 0.000 |
| 7 | Left | Vis | Visual | 0.328 | 0.298 | 0.239 | 0.005 | 0.000 |
| 8 | Left | Vis | Visual | 0.365 | 0.254 | 0.861 | 0.005 | 0.000 |
| 9 | Left | Vis | Visual | 0.209 | 0.173 | 0.345 | 0.006 | 0.000 |
| 10 | Left | SomMot | Somatomotor | 0.364 | 0.315 | 0.430 | 0.005 | 0.000 |
| 11 | Left | SomMot | Somatomotor | 0.054 | 0.053 | 0.026 | 0.007 | 0.000 |
| 12 | Left | SomMot | Somatomotor | 0.050 | 0.041 | 0.115 | 0.008 | 0.000 |
| 13 | Left | SomMot | Somatomotor | 0.016 | 0.013 | 0.034 | 0.008 | 0.000 |
| 16 | Left | DorsAttn | Posterior | 0.199 | 0.191 | 0.091 | 0.006 | 0.000 |
| 17 | Left | DorsAttn | Posterior | 0.148 | 0.160 | -0.118 | 0.006 | 0.000 |
| 18 | Left | DorsAttn | Posterior | 0.278 | 0.176 | 0.834 | 0.005 | 0.000 |
| 19 | Left | DorsAttn | Posterior | 0.132 | 0.125 | 0.068 | 0.006 | 0.000 |
| 20 | Left | DorsAttn | Posterior | 0.175 | 0.112 | 0.604 | 0.006 | 0.000 |
| 21 | Left | DorsAttn | Posterior | 0.174 | 0.162 | 0.113 | 0.006 | 0.000 |
| 22 | Left | DorsAttn | Precentral Ventral | 0.080 | 0.115 | -0.411 | 0.007 | 0.000 |
| 23 | Left | DorsAttn | Frontal Eye Fields | 0.121 | 0.102 | 0.226 | 0.006 | 0.000 |
| 24 | Left | SalVentAttn | Parietal Operculum | 0.087 | 0.116 | -0.322 | 0.007 | 0.000 |
| 25 | Left | SalVentAttn | Frontal Operculum/Insula | 0.040 | 0.060 | -0.308 | 0.008 | 0.000 |
| 26 | Left | SalVentAttn | Frontal Operculum/Insula | 0.026 | 0.037 | -0.152 | 0.008 | 0.000 |
| 27 | Left | SalVentAttn | Lateral Prefrontal Cortex | 0.047 | 0.044 | 0.052 | 0.008 | 0.000 |
| 28 | Left | SalVentAttn | Medial | 0.023 | 0.030 | -0.101 | 0.008 | 0.000 |
| 29 | Left | SalVentAttn | Medial | 0.112 | 0.089 | 0.277 | 0.007 | 0.000 |
| 30 | Left | SalVentAttn | Medial | 0.050 | 0.037 | 0.184 | 0.008 | 0.000 |
| 31 | Left | Limbic | Orbitofrontal Cortex | 0.025 | 0.017 | 0.165 | 0.008 | 0.000 |
| 32 | Left | Limbic | Temporal Pole | 0.090 | 0.079 | 0.154 | 0.007 | 0.000 |
| 33 | Left | Limbic | Temporal Pole | 0.085 | 0.046 | 0.489 | 0.007 | 0.000 |

| | | | | | | | | |
|---|---|---|---|---|---|---|---|---|
| 34 | Left | Cont | Parietal | 0.111 | 0.095 | 0.186 | 0.007 | 0.000 |
| 35 | Left | Cont | Lateral Prefrontal Cortex | 0.063 | 0.049 | 0.174 | 0.008 | 0.000 |
| 37 | Left | Cont | Cingulate Cortex | 0.067 | 0.063 | 0.064 | 0.008 | 0.000 |
| 38 | Left | Default | Temporal | 0.229 | 0.165 | 0.626 | 0.006 | 0.000 |
| 39 | Left | Default | Temporal | 0.260 | 0.176 | 0.744 | 0.006 | 0.000 |
| 40 | Left | Default | Parietal | 0.199 | 0.177 | 0.230 | 0.007 | 0.000 |
| 41 | Left | Default | Parietal | 0.106 | 0.074 | 0.365 | 0.007 | 0.000 |
| 42 | Left | Default | Prefrontal Cortex | 0.058 | 0.043 | 0.224 | 0.008 | 0.000 |
| 43 | Left | Default | Prefrontal Cortex | 0.114 | 0.056 | 0.610 | 0.007 | 0.000 |
| 45 | Left | Default | Prefrontal Cortex | 0.062 | 0.045 | 0.218 | 0.008 | 0.000 |
| 46 | Left | Default | Prefrontal Cortex | 0.101 | 0.049 | 0.576 | 0.007 | 0.000 |
| 47 | Left | Default | Prefrontal Cortex | 0.095 | 0.054 | 0.477 | 0.007 | 0.000 |
| 48 | Left | Default | Prefrontal Cortex | 0.081 | 0.042 | 0.464 | 0.007 | 0.000 |
| 49 | Left | Default | Precuneus/Posterior Cingulate Cortex | 0.141 | 0.115 | 0.293 | 0.006 | 0.000 |
| 50 | Left | Default | Precuneus/Posterior Cingulate Cortex | 0.107 | 0.084 | 0.268 | 0.007 | 0.000 |
| 51 | Right | Vis | Visual | 0.247 | 0.165 | 0.794 | 0.005 | 0.000 |
| 52 | Right | Vis | Visual | 0.360 | 0.271 | 0.704 | 0.005 | 0.000 |
| 53 | Right | Vis | Visual | 0.275 | 0.221 | 0.506 | 0.005 | 0.000 |
| 54 | Right | Vis | Visual | 0.231 | 0.278 | -0.395 | 0.005 | 0.000 |
| 55 | Right | Vis | Visual | 0.268 | 0.225 | 0.363 | 0.005 | 0.000 |
| 56 | Right | Vis | Visual | 0.228 | 0.187 | 0.386 | 0.005 | 0.000 |
| 57 | Right | Vis | Visual | 0.378 | 0.251 | 0.955 | 0.005 | 0.000 |
| 58 | Right | Vis | Visual | 0.275 | 0.209 | 0.525 | 0.005 | 0.000 |
| 59 | Right | SomMot | Somatomotor | 0.317 | 0.280 | 0.329 | 0.005 | 0.000 |
| 60 | Right | SomMot | Somatomotor | 0.026 | 0.024 | 0.032 | 0.008 | 0.000 |
| 61 | Right | SomMot | Somatomotor | 0.052 | 0.034 | 0.251 | 0.008 | 0.000 |
| 63 | Right | SomMot | Somatomotor | 0.023 | 0.021 | 0.020 | 0.008 | 0.020 |
| 64 | Right | SomMot | Somatomotor | 0.020 | 0.024 | -0.051 | 0.008 | 0.000 |
| 65 | Right | SomMot | Somatomotor | 0.091 | 0.063 | 0.288 | 0.007 | 0.000 |
| 66 | Right | SomMot | Somatomotor | 0.032 | 0.034 | -0.027 | 0.008 | 0.001 |
| 67 | Right | DorsAttn | Posterior | 0.266 | 0.200 | 0.591 | 0.005 | 0.000 |
| 68 | Right | DorsAttn | Posterior | 0.177 | 0.142 | 0.337 | 0.006 | 0.000 |
| 69 | Right | DorsAttn | Posterior | 0.190 | 0.161 | 0.288 | 0.006 | 0.000 |
| 70 | Right | DorsAttn | Posterior | 0.280 | 0.161 | 0.923 | 0.005 | 0.000 |
| 71 | Right | DorsAttn | Posterior | 0.190 | 0.131 | 0.520 | 0.006 | 0.000 |

| | | | | | | | | |
|---|---|---|---|---|---|---|---|---|
| 72 | Right | DorsAttn | Precentral Ventral | 0.131 | 0.118 | 0.146 | 0.007 | 0.000 |
| 73 | Right | DorsAttn | Frontal Eye Fields | 0.177 | 0.116 | 0.594 | 0.006 | 0.000 |
| 74 | Right | SalVentAttn | Temporal Occipital Parietal | 0.254 | 0.202 | 0.483 | 0.006 | 0.000 |
| 75 | Right | SalVentAttn | Temporal Occipital Parietal | 0.120 | 0.133 | -0.125 | 0.006 | 0.000 |
| 76 | Right | SalVentAttn | Frontal Operculum/Insula | 0.033 | 0.043 | -0.135 | 0.008 | 0.000 |
| 77 | Right | SalVentAttn | Medial | 0.102 | 0.075 | 0.331 | 0.007 | 0.000 |
| 79 | Right | Limbic | Orbitofrontal Cortex | 0.020 | 0.014 | 0.132 | 0.008 | 0.000 |
| 80 | Right | Limbic | Temporal Pole | 0.097 | 0.076 | 0.311 | 0.007 | 0.000 |
| 81 | Right | Cont | Parietal | 0.093 | 0.109 | -0.162 | 0.007 | 0.000 |
| 82 | Right | Cont | Parietal | 0.096 | 0.083 | 0.149 | 0.007 | 0.000 |
| 84 | Right | Cont | Lateral Prefrontal Cortex | 0.079 | 0.064 | 0.181 | 0.007 | 0.000 |
| 85 | Right | Cont | Lateral Prefrontal Cortex | 0.056 | 0.061 | -0.056 | 0.008 | 0.000 |
| 86 | Right | Cont | Lateral Prefrontal Cortex | 0.062 | 0.073 | -0.130 | 0.008 | 0.000 |
| 87 | Right | Cont | Cingulate Cortex | 0.065 | 0.069 | -0.055 | 0.008 | 0.000 |
| 88 | Right | Cont | Medial Posterior Prefrontal Cortex | 0.028 | 0.042 | -0.182 | 0.008 | 0.000 |
| 89 | Right | Cont | Precuneus | 0.129 | 0.119 | 0.116 | 0.007 | 0.000 |
| 90 | Right | Default | Parietal | 0.089 | 0.102 | -0.142 | 0.007 | 0.000 |
| 91 | Right | Default | Temporal | 0.065 | 0.056 | 0.131 | 0.008 | 0.000 |
| 92 | Right | Default | Temporal | 0.224 | 0.188 | 0.391 | 0.006 | 0.000 |
| 93 | Right | Default | Temporal | 0.248 | 0.205 | 0.410 | 0.006 | 0.000 |
| 94 | Right | Default | Temporal | 0.030 | 0.039 | -0.132 | 0.008 | 0.000 |
| 95 | Right | Default | Ventral Prefrontal Cortex | 0.093 | 0.088 | 0.062 | 0.007 | 0.000 |
| 96 | Right | Default | Ventral Prefrontal Cortex | 0.051 | 0.041 | 0.137 | 0.008 | 0.000 |
| 97 | Right | Default | Dorsomedial Prefrontal Cortex | 0.106 | 0.067 | 0.454 | 0.007 | 0.000 |
| 98 | Right | Default | Dorsomedial Prefrontal Cortex | 0.095 | 0.066 | 0.348 | 0.007 | 0.000 |
| 99 | Right | Default | Precuneus/Posterior Cingulate Cortex | 0.177 | 0.148 | 0.304 | 0.006 | 0.000 |
| 100 | Left | Default | Precuneus/Posterior Cingulate Cortex | 0.105 | 0.094 | 0.135 | 0.007 | 0.000 |

Subcortical Results

| Parcel No. | Hemisphere | Region | $ISC_{social}$ | $ISC_{nonsocial}$ | β | *SE* | *p* |
|---|---|---|---|---|---|---|---|
| 103 | Left | Putamen | 0.021 | 0.017 | 0.076 | 0.008 | 0.000 |
| 104 | Left | Pallidum | 0.005 | 0.004 | 0.030 | 0.008 | 0.000 |
| 105 | Left | Hippocampus | 0.038 | 0.037 | 0.036 | 0.008 | 0.000 |
| 106 | Left | Amygdala | 0.046 | 0.043 | 0.070 | 0.008 | 0.000 |
| 107 | Left | Accumbens | 0.008 | 0.010 | -0.034 | 0.008 | 0.000 |
| 108 | Left | Thalamus | 0.035 | 0.032 | 0.058 | 0.008 | 0.000 |

| | | | | | | | |
|---|---|---|---|---|---|---|---|
| 109 | Right | Caudate | 0.025 | 0.028 | -0.054 | 0.008 | 0.000 |
| 110 | Right | Putamen | 0.020 | 0.019 | 0.028 | 0.008 | 0.001 |
| 113 | Right | Amygdala | 0.057 | 0.049 | 0.151 | 0.008 | 0.000 |
| 114 | Right | Accumbens | 0.009 | 0.012 | -0.053 | 0.008 | 0.000 |

*Note.* Parcel numbers for cortical regions correspond to the Schaefer 100 cortical parcellation (Schaefer et al., 2018), whereas parcel numbers for subcortical regions are arbitrary numbers assigned by researchers. Component names are from the labels provided by Yeo et al., (2011). Although linear mixed effects modelling considers the fully crossed random effects in comparing differences in the ISCs for each context, we provide mean ISCs in each context to enhance interpretability and facilitate direct comparison. **β** indicates standardized coefficients analyzed with the contrast of $ISC_{social} > ISC_{nonsocial}$. *SE* indicates standard error, and *p*-values were corrected for multiple comparisons using a false discovery rate at $p < .05$. Vis = Visual, SomMot = Somatomotor, DorsAttn = Dorsal Attention, SalVentAttn = Salience/Ventral Attention, Cont = Control.

**Table S4.** Brain regions showing significant main effect of context when controlling for covariates (demographic similarities, kinship, depression, and objective levels of social connection).

| | | | Cortical Results | | | | | |
|---|---|---|---|---|---|---|---|---|
| Parcel No. | Hemisphere | Network | Component | $ISC_{social}$ | $ISC_{nonsocial}$ | β | *SE* | *p* |
| 1 | Left | Vis | Visual | 0.229 | 0.173 | 0.597 | 0.005 | 0.000 |
| 2 | Left | Vis | Visual | 0.300 | 0.264 | 0.303 | 0.005 | 0.000 |
| 3 | Left | Vis | Visual | 0.311 | 0.229 | 0.669 | 0.005 | 0.000 |
| 4 | Left | Vis | Visual | 0.219 | 0.269 | -0.423 | 0.005 | 0.000 |
| 5 | Left | Vis | Visual | 0.305 | 0.325 | -0.149 | 0.005 | 0.000 |
| 6 | Left | Vis | Visual | 0.241 | 0.182 | 0.530 | 0.005 | 0.000 |
| 7 | Left | Vis | Visual | 0.328 | 0.298 | 0.239 | 0.005 | 0.000 |
| 8 | Left | Vis | Visual | 0.365 | 0.254 | 0.861 | 0.005 | 0.000 |
| 9 | Left | Vis | Visual | 0.209 | 0.173 | 0.345 | 0.006 | 0.000 |
| 10 | Left | SomMot | Somatomotor | 0.364 | 0.315 | 0.430 | 0.005 | 0.000 |
| 11 | Left | SomMot | Somatomotor | 0.054 | 0.053 | 0.026 | 0.007 | 0.000 |
| 12 | Left | SomMot | Somatomotor | 0.050 | 0.041 | 0.115 | 0.008 | 0.000 |
| 13 | Left | SomMot | Somatomotor | 0.016 | 0.013 | 0.034 | 0.008 | 0.000 |
| 16 | Left | DorsAttn | Posterior | 0.199 | 0.191 | 0.091 | 0.006 | 0.000 |
| 17 | Left | DorsAttn | Posterior | 0.148 | 0.160 | -0.118 | 0.006 | 0.000 |
| 18 | Left | DorsAttn | Posterior | 0.278 | 0.176 | 0.834 | 0.005 | 0.000 |
| 19 | Left | DorsAttn | Posterior | 0.132 | 0.125 | 0.068 | 0.006 | 0.000 |
| 20 | Left | DorsAttn | Posterior | 0.175 | 0.112 | 0.604 | 0.006 | 0.000 |
| 21 | Left | DorsAttn | Posterior | 0.174 | 0.162 | 0.113 | 0.006 | 0.000 |
| 22 | Left | DorsAttn | Precentral Ventral | 0.080 | 0.115 | -0.411 | 0.007 | 0.000 |
| 23 | Left | DorsAttn | Frontal Eye Fields | 0.121 | 0.102 | 0.226 | 0.006 | 0.000 |
| 24 | Left | SalVentAttn | Parietal Operculum | 0.087 | 0.116 | -0.322 | 0.007 | 0.000 |
| 25 | Left | SalVentAttn | Frontal Operculum/Insula | 0.040 | 0.060 | -0.308 | 0.008 | 0.000 |
| 26 | Left | SalVentAttn | Frontal Operculum/Insula | 0.026 | 0.037 | -0.152 | 0.008 | 0.000 |
| 27 | Left | SalVentAttn | Lateral Prefrontal Cortex | 0.047 | 0.044 | 0.052 | 0.008 | 0.000 |
| 28 | Left | SalVentAttn | Medial | 0.023 | 0.030 | -0.101 | 0.008 | 0.000 |
| 29 | Left | SalVentAttn | Medial | 0.112 | 0.089 | 0.277 | 0.007 | 0.000 |
| 30 | Left | SalVentAttn | Medial | 0.050 | 0.037 | 0.184 | 0.008 | 0.000 |
| 31 | Left | Limbic | Orbitofrontal Cortex | 0.025 | 0.017 | 0.165 | 0.008 | 0.000 |
| 32 | Left | Limbic | Temporal Pole | 0.090 | 0.079 | 0.154 | 0.007 | 0.000 |

| | | | | | | | | |
|---|---|---|---|---|---|---|---|---|
| 33 | Left | Limbic | Temporal Pole | 0.085 | 0.046 | 0.489 | 0.007 | 0.000 |
| 34 | Left | Cont | Parietal | 0.111 | 0.095 | 0.186 | 0.007 | 0.000 |
| 35 | Left | Cont | Lateral Prefrontal Cortex | 0.063 | 0.049 | 0.174 | 0.008 | 0.000 |
| 37 | Left | Cont | Cingulate Cortex | 0.067 | 0.063 | 0.064 | 0.008 | 0.000 |
| 38 | Left | Default | Temporal | 0.229 | 0.165 | 0.626 | 0.006 | 0.000 |
| 39 | Left | Default | Temporal | 0.260 | 0.176 | 0.744 | 0.006 | 0.000 |
| 40 | Left | Default | Parietal | 0.199 | 0.177 | 0.230 | 0.007 | 0.000 |
| 41 | Left | Default | Parietal | 0.106 | 0.074 | 0.365 | 0.007 | 0.000 |
| 42 | Left | Default | Prefrontal Cortex | 0.058 | 0.043 | 0.224 | 0.008 | 0.000 |
| 43 | Left | Default | Prefrontal Cortex | 0.114 | 0.056 | 0.610 | 0.007 | 0.000 |
| 45 | Left | Default | Prefrontal Cortex | 0.062 | 0.045 | 0.218 | 0.008 | 0.000 |
| 46 | Left | Default | Prefrontal Cortex | 0.101 | 0.049 | 0.576 | 0.007 | 0.000 |
| 47 | Left | Default | Prefrontal Cortex | 0.095 | 0.054 | 0.477 | 0.007 | 0.000 |
| 48 | Left | Default | Prefrontal Cortex | 0.081 | 0.042 | 0.464 | 0.007 | 0.000 |
| 49 | Left | Default | Precuneus/Posterior Cingulate Cortex | 0.141 | 0.115 | 0.293 | 0.006 | 0.000 |
| 50 | Left | Default | Precuneus/Posterior Cingulate Cortex | 0.107 | 0.084 | 0.268 | 0.007 | 0.000 |
| 51 | Right | Vis | Visual | 0.247 | 0.165 | 0.794 | 0.005 | 0.000 |
| 52 | Right | Vis | Visual | 0.360 | 0.271 | 0.704 | 0.005 | 0.000 |
| 53 | Right | Vis | Visual | 0.275 | 0.221 | 0.506 | 0.005 | 0.000 |
| 54 | Right | Vis | Visual | 0.231 | 0.278 | -0.395 | 0.005 | 0.000 |
| 55 | Right | Vis | Visual | 0.268 | 0.225 | 0.363 | 0.005 | 0.000 |
| 56 | Right | Vis | Visual | 0.228 | 0.187 | 0.386 | 0.005 | 0.000 |
| 57 | Right | Vis | Visual | 0.378 | 0.251 | 0.955 | 0.005 | 0.000 |
| 58 | Right | Vis | Visual | 0.275 | 0.209 | 0.525 | 0.005 | 0.000 |
| 59 | Right | SomMot | Somatomotor | 0.317 | 0.280 | 0.329 | 0.005 | 0.000 |
| 60 | Right | SomMot | Somatomotor | 0.026 | 0.024 | 0.032 | 0.008 | 0.000 |
| 61 | Right | SomMot | Somatomotor | 0.052 | 0.034 | 0.251 | 0.008 | 0.000 |
| 63 | Right | SomMot | Somatomotor | 0.023 | 0.021 | 0.020 | 0.008 | 0.020 |
| 64 | Right | SomMot | Somatomotor | 0.020 | 0.024 | -0.051 | 0.008 | 0.000 |
| 65 | Right | SomMot | Somatomotor | 0.091 | 0.063 | 0.288 | 0.007 | 0.000 |
| 66 | Right | SomMot | Somatomotor | 0.032 | 0.034 | -0.027 | 0.008 | 0.001 |
| 67 | Right | DorsAttn | Posterior | 0.266 | 0.200 | 0.591 | 0.005 | 0.000 |
| 68 | Right | DorsAttn | Posterior | 0.177 | 0.142 | 0.337 | 0.006 | 0.000 |
| 69 | Right | DorsAttn | Posterior | 0.190 | 0.161 | 0.288 | 0.006 | 0.000 |
| 70 | Right | DorsAttn | Posterior | 0.280 | 0.161 | 0.923 | 0.005 | 0.000 |

| | | | | | | | | |
|---|---|---|---|---|---|---|---|---|
| 71 | Right | DorsAttn | Posterior | 0.190 | 0.131 | 0.520 | 0.006 | 0.000 |
| 72 | Right | DorsAttn | Precentral Ventral | 0.131 | 0.118 | 0.146 | 0.007 | 0.000 |
| 73 | Right | DorsAttn | Frontal Eye Fields | 0.177 | 0.116 | 0.594 | 0.006 | 0.000 |
| 74 | Right | SalVentAttn | Temporal Occipital Parietal | 0.254 | 0.202 | 0.483 | 0.006 | 0.000 |
| 75 | Right | SalVentAttn | Temporal Occipital Parietal | 0.120 | 0.133 | -0.125 | 0.006 | 0.000 |
| 76 | Right | SalVentAttn | Frontal Operculum/Insula | 0.033 | 0.043 | -0.135 | 0.008 | 0.000 |
| 77 | Right | SalVentAttn | Medial | 0.102 | 0.075 | 0.331 | 0.007 | 0.000 |
| 79 | Right | Limbic | Orbitofrontal Cortex | 0.020 | 0.014 | 0.132 | 0.008 | 0.000 |
| 80 | Right | Limbic | Temporal Pole | 0.097 | 0.076 | 0.311 | 0.007 | 0.000 |
| 81 | Right | Cont | Parietal | 0.093 | 0.109 | -0.162 | 0.007 | 0.000 |
| 82 | Right | Cont | Parietal | 0.096 | 0.083 | 0.149 | 0.007 | 0.000 |
| 84 | Right | Cont | Lateral Prefrontal Cortex | 0.079 | 0.064 | 0.181 | 0.007 | 0.000 |
| 85 | Right | Cont | Lateral Prefrontal Cortex | 0.056 | 0.061 | -0.056 | 0.008 | 0.000 |
| 86 | Right | Cont | Lateral Prefrontal Cortex | 0.062 | 0.073 | -0.130 | 0.008 | 0.000 |
| 87 | Right | Cont | Cingulate Cortex | 0.065 | 0.069 | -0.055 | 0.008 | 0.000 |
| 88 | Right | Cont | Medial Posterior Prefrontal Cortex | 0.028 | 0.042 | -0.182 | 0.008 | 0.000 |
| 89 | Right | Cont | Precuneus | 0.129 | 0.119 | 0.116 | 0.007 | 0.000 |
| 90 | Right | Default | Parietal | 0.089 | 0.102 | -0.142 | 0.007 | 0.000 |
| 91 | Right | Default | Temporal | 0.065 | 0.056 | 0.131 | 0.008 | 0.000 |
| 92 | Right | Default | Temporal | 0.224 | 0.188 | 0.391 | 0.006 | 0.000 |
| 93 | Right | Default | Temporal | 0.248 | 0.205 | 0.410 | 0.006 | 0.000 |
| 94 | Right | Default | Temporal | 0.030 | 0.039 | -0.132 | 0.008 | 0.000 |
| 95 | Right | Default | Ventral Prefrontal Cortex | 0.093 | 0.088 | 0.062 | 0.007 | 0.000 |
| 96 | Right | Default | Ventral Prefrontal Cortex | 0.051 | 0.041 | 0.137 | 0.008 | 0.000 |
| 97 | Right | Default | Dorsomedial Prefrontal Cortex | 0.106 | 0.067 | 0.454 | 0.007 | 0.000 |
| 98 | Right | Default | Dorsomedial Prefrontal Cortex | 0.095 | 0.066 | 0.348 | 0.007 | 0.000 |
| 99 | Right | Default | Precuneus/Posterior Cingulate Cortex | 0.177 | 0.148 | 0.304 | 0.006 | 0.000 |
| 100 | Left | Default | Precuneus/Posterior Cingulate Cortex | 0.105 | 0.094 | 0.135 | 0.007 | 0.000 |

Subcortical Results

| Parcel No. | Hemisphere | Region | $ISC_{social}$ | $ISC_{nonsocial}$ | $\beta$ | *SE* | *p* |
|---|---|---|---|---|---|---|---|
| 103 | Left | Putamen | 0.021 | 0.017 | 0.076 | 0.008 | 0.000 |
| 104 | Left | Pallidum | 0.005 | 0.004 | 0.030 | 0.008 | 0.000 |
| 105 | Left | Hippocampus | 0.038 | 0.037 | 0.036 | 0.008 | 0.000 |
| 106 | Left | Amygdala | 0.046 | 0.043 | 0.070 | 0.008 | 0.000 |
| 107 | Left | Accumbens | 0.008 | 0.010 | -0.034 | 0.008 | 0.000 |

| | | | | | | | |
|---|---|---|---|---|---|---|---|
| 108 | Left | Thalamus | 0.035 | 0.032 | 0.058 | 0.008 | 0.000 |
| 109 | Right | Caudate | 0.025 | 0.028 | -0.054 | 0.008 | 0.000 |
| 110 | Right | Putamen | 0.020 | 0.019 | 0.028 | 0.008 | 0.001 |
| 113 | Right | Amygdala | 0.057 | 0.049 | 0.151 | 0.008 | 0.000 |
| 114 | Right | Accumbens | 0.009 | 0.012 | -0.053 | 0.008 | 0.000 |

*Note.* Parcel numbers for cortical regions correspond to the Schaefer 100 cortical parcellation (Schaefer et al., 2018), whereas parcel numbers for subcortical regions are arbitrary numbers assigned by researchers. Component names are from the labels provided by Yeo et al., (2011). Although linear mixed effects modelling considers the fully crossed random effects in comparing differences in the ISCs for each context, we provide mean ISCs in each context to enhance interpretability and facilitate direct comparison. **β** indicates standardized coefficients analyzed with the contrast of $ISC_{social} > ISC_{nonsocial}$. *SE* indicates standard error, and *p*-values were corrected for multiple comparisons using a false discovery rate at $p < .05$. Vis = Visual, SomMot = Somatomotor, DorsAttn = Dorsal Attention, SalVentAttn = Salience/Ventral Attention, Cont = Control.

**Table S5.** Brain regions showing significant interaction effects between loneliness and context ($ISC_{hh}$ > $ISC_{ll}$).

| Cortical Results | | | | | | | | | | |
|---|---|---|---|---|---|---|---|---|---|---|
| Parcel No. | Hemisphere | Network | Component | $ISC_{hh.s}$ | $ISC_{ll.s}$ | $ISC_{hh.ns}$ | $ISC_{ll.ns}$ | β | *SE* | *p* |
| 1 | Left | Vis | Visual | 0.224 | 0.234 | 0.165 | 0.180 | 0.037 | 0.015 | 0.015 |
| 2 | Left | Vis | Visual | 0.291 | 0.310 | 0.261 | 0.268 | -0.098 | 0.014 | 0.000 |
| 3 | Left | Vis | Visual | 0.308 | 0.315 | 0.221 | 0.237 | 0.074 | 0.013 | 0.000 |
| 4 | Left | Vis | Visual | 0.206 | 0.231 | 0.265 | 0.274 | -0.121 | 0.014 | 0.000 |
| 5 | Left | Vis | Visual | 0.291 | 0.319 | 0.317 | 0.332 | -0.097 | 0.013 | 0.000 |
| 6 | Left | Vis | Visual | 0.234 | 0.248 | 0.179 | 0.183 | -0.088 | 0.014 | 0.000 |
| 7 | Left | Vis | Visual | 0.318 | 0.337 | 0.294 | 0.303 | -0.081 | 0.013 | 0.000 |
| 8 | Left | Vis | Visual | 0.361 | 0.369 | 0.246 | 0.262 | 0.062 | 0.013 | 0.000 |
| 9 | Left | Vis | Visual | 0.212 | 0.208 | 0.166 | 0.178 | 0.141 | 0.015 | 0.000 |
| 10 | Left | SomMot | Somatomotor | 0.353 | 0.375 | 0.320 | 0.311 | -0.258 | 0.013 | 0.000 |
| 11 | Left | SomMot | Somatomotor | 0.054 | 0.056 | 0.057 | 0.048 | -0.171 | 0.020 | 0.000 |
| 12 | Left | SomMot | Somatomotor | 0.050 | 0.050 | 0.045 | 0.038 | -0.100 | 0.021 | 0.000 |
| 13 | Left | SomMot | Somatomotor | 0.015 | 0.016 | 0.018 | 0.008 | -0.143 | 0.022 | 0.000 |
| 14 | Left | SomMot | Somatomotor | 0.021 | 0.022 | 0.027 | 0.018 | -0.118 | 0.022 | 0.000 |
| 15 | Left | SomMot | Somatomotor | 0.030 | 0.031 | 0.036 | 0.026 | -0.154 | 0.022 | 0.000 |
| 16 | Left | DorsAttn | Posterior | 0.195 | 0.203 | 0.188 | 0.193 | -0.034 | 0.017 | 0.050 |
| 17 | Left | DorsAttn | Posterior | 0.151 | 0.147 | 0.157 | 0.163 | 0.102 | 0.016 | 0.000 |
| 18 | Left | DorsAttn | Posterior | 0.265 | 0.290 | 0.169 | 0.183 | -0.097 | 0.014 | 0.000 |
| 19 | Left | DorsAttn | Posterior | 0.132 | 0.133 | 0.123 | 0.128 | 0.040 | 0.016 | 0.016 |
| 21 | Left | DorsAttn | Posterior | 0.168 | 0.181 | 0.162 | 0.163 | -0.096 | 0.015 | 0.000 |
| 24 | Left | SalVentAttn | Parietal Operculum | 0.086 | 0.090 | 0.117 | 0.115 | -0.072 | 0.018 | 0.000 |
| 26 | Left | SalVentAttn | Frontal Operculum/Insula | 0.028 | 0.026 | 0.034 | 0.037 | 0.078 | 0.022 | 0.001 |
| 27 | Left | SalVentAttn | Lateral Prefrontal Cortex | 0.050 | 0.047 | 0.040 | 0.047 | 0.118 | 0.021 | 0.000 |
| 28 | Left | SalVentAttn | Medial | 0.023 | 0.022 | 0.026 | 0.031 | 0.097 | 0.022 | 0.000 |
| 29 | Left | SalVentAttn | Medial | 0.116 | 0.109 | 0.089 | 0.088 | 0.068 | 0.018 | 0.000 |
| 31 | Left | Limbic | Orbitofrontal Cortex | 0.024 | 0.027 | 0.014 | 0.020 | 0.079 | 0.022 | 0.001 |
| 32 | Left | Limbic | Temporal Pole | 0.085 | 0.096 | 0.078 | 0.080 | -0.124 | 0.019 | 0.000 |
| 33 | Left | Limbic | Temporal Pole | 0.078 | 0.090 | 0.044 | 0.049 | -0.105 | 0.019 | 0.000 |
| 34 | Left | Cont | Parietal | 0.105 | 0.116 | 0.086 | 0.103 | 0.060 | 0.019 | 0.002 |
| 35 | Left | Cont | Lateral Prefrontal Cortex | 0.063 | 0.063 | 0.043 | 0.055 | 0.151 | 0.021 | 0.000 |
| 36 | Left | Cont | Precuneus | 0.101 | 0.100 | 0.094 | 0.108 | 0.185 | 0.020 | 0.000 |

| | | | | | | | | | | |
|---|---|---|---|---|---|---|---|---|---|---|
| 37 | Left | Cont | Cingulate Cortex | 0.067 | 0.067 | 0.059 | 0.065 | 0.072 | 0.021 | 0.001 |
| 38 | Left | Default | Temporal | 0.215 | 0.245 | 0.162 | 0.167 | -0.237 | 0.016 | 0.000 |
| 39 | Left | Default | Temporal | 0.240 | 0.279 | 0.175 | 0.177 | -0.329 | 0.015 | 0.000 |
| 40 | Left | Default | Parietal | 0.188 | 0.211 | 0.173 | 0.180 | -0.154 | 0.018 | 0.000 |
| 42 | Left | Default | Prefrontal Cortex | 0.053 | 0.063 | 0.039 | 0.045 | -0.048 | 0.020 | 0.026 |
| 43 | Left | Default | Prefrontal Cortex | 0.112 | 0.114 | 0.050 | 0.060 | 0.085 | 0.018 | 0.000 |
| 44 | Left | Default | Prefrontal Cortex | 0.034 | 0.038 | 0.035 | 0.035 | -0.050 | 0.022 | 0.027 |
| 45 | Left | Default | Prefrontal Cortex | 0.060 | 0.063 | 0.041 | 0.048 | 0.048 | 0.021 | 0.027 |
| 46 | Left | Default | Prefrontal Cortex | 0.091 | 0.110 | 0.046 | 0.051 | -0.171 | 0.019 | 0.000 |
| 50 | Left | Default | Precuneus/Posterior Cingulate Cortex | 0.098 | 0.115 | 0.079 | 0.089 | -0.086 | 0.019 | 0.000 |
| 54 | Right | Vis | Visual | 0.220 | 0.242 | 0.273 | 0.285 | -0.073 | 0.014 | 0.000 |
| 55 | Right | Vis | Visual | 0.261 | 0.275 | 0.214 | 0.235 | 0.050 | 0.014 | 0.001 |
| 56 | Right | Vis | Visual | 0.231 | 0.226 | 0.183 | 0.189 | 0.097 | 0.014 | 0.000 |
| 58 | Right | Vis | Visual | 0.275 | 0.275 | 0.198 | 0.218 | 0.158 | 0.014 | 0.000 |
| 59 | Right | SomMot | Somatomotor | 0.309 | 0.325 | 0.285 | 0.276 | -0.205 | 0.013 | 0.000 |
| 60 | Right | SomMot | Somatomotor | 0.027 | 0.026 | 0.029 | 0.020 | -0.136 | 0.022 | 0.000 |
| 61 | Right | SomMot | Somatomotor | 0.054 | 0.051 | 0.038 | 0.030 | -0.076 | 0.021 | 0.001 |
| 62 | Right | SomMot | Somatomotor | 0.013 | 0.013 | 0.016 | 0.007 | -0.127 | 0.022 | 0.000 |
| 63 | Right | SomMot | Somatomotor | 0.023 | 0.023 | 0.023 | 0.019 | -0.065 | 0.022 | 0.005 |
| 64 | Right | SomMot | Somatomotor | 0.021 | 0.019 | 0.030 | 0.019 | -0.124 | 0.022 | 0.000 |
| 65 | Right | SomMot | Somatomotor | 0.088 | 0.093 | 0.065 | 0.062 | -0.090 | 0.018 | 0.000 |
| 66 | Right | SomMot | Somatomotor | 0.033 | 0.033 | 0.041 | 0.030 | -0.144 | 0.022 | 0.000 |
| 67 | Right | DorsAttn | Posterior | 0.251 | 0.282 | 0.189 | 0.210 | -0.083 | 0.015 | 0.000 |
| 70 | Right | DorsAttn | Posterior | 0.268 | 0.291 | 0.152 | 0.168 | -0.067 | 0.014 | 0.000 |
| 71 | Right | DorsAttn | Posterior | 0.180 | 0.199 | 0.130 | 0.132 | -0.141 | 0.015 | 0.000 |
| 73 | Right | DorsAttn | Frontal Eye Fields | 0.174 | 0.179 | 0.114 | 0.116 | -0.032 | 0.015 | 0.045 |
| 74 | Right | SalVentAttn | Temporal Occipital Parietal | 0.242 | 0.265 | 0.205 | 0.199 | -0.268 | 0.015 | 0.000 |
| 75 | Right | SalVentAttn | Temporal Occipital Parietal | 0.123 | 0.120 | 0.132 | 0.133 | 0.045 | 0.017 | 0.010 |
| 76 | Right | SalVentAttn | Frontal Operculum/Insula | 0.036 | 0.032 | 0.041 | 0.044 | 0.100 | 0.022 | 0.000 |
| 79 | Right | Limbic | Orbitofrontal Cortex | 0.021 | 0.019 | 0.011 | 0.017 | 0.160 | 0.022 | 0.000 |
| 80 | Right | Limbic | Temporal Pole | 0.090 | 0.106 | 0.076 | 0.076 | -0.238 | 0.019 | 0.000 |
| 81 | Right | Cont | Parietal | 0.091 | 0.097 | 0.102 | 0.113 | 0.050 | 0.019 | 0.010 |
| 82 | Right | Cont | Parietal | 0.087 | 0.105 | 0.079 | 0.086 | -0.121 | 0.020 | 0.000 |
| 83 | Right | Cont | Lateral Prefrontal Cortex | 0.053 | 0.066 | 0.059 | 0.059 | -0.162 | 0.021 | 0.000 |
| 84 | Right | Cont | Lateral Prefrontal Cortex | 0.074 | 0.085 | 0.061 | 0.065 | -0.067 | 0.020 | 0.001 |

| | | | | | | | | | | |
|---|---|---|---|---|---|---|---|---|---|---|
| 85 | Right | Cont | Lateral Prefrontal Cortex | 0.057 | 0.057 | 0.057 | 0.064 | 0.076 | 0.021 | 0.000 |
| 86 | Right | Cont | Lateral Prefrontal Cortex | 0.051 | 0.074 | 0.069 | 0.076 | -0.195 | 0.020 | 0.000 |
| 87 | Right | Cont | Cingulate Cortex | 0.066 | 0.064 | 0.067 | 0.070 | 0.078 | 0.021 | 0.000 |
| 88 | Right | Cont | Medial Posterior Prefrontal Cortex | 0.027 | 0.030 | 0.035 | 0.045 | 0.095 | 0.021 | 0.000 |
| 89 | Right | Cont | Precuneus | 0.129 | 0.130 | 0.116 | 0.122 | 0.050 | 0.018 | 0.010 |
| 90 | Right | Default | Parietal | 0.080 | 0.098 | 0.097 | 0.105 | -0.112 | 0.019 | 0.000 |
| 91 | Right | Default | Temporal | 0.062 | 0.068 | 0.054 | 0.057 | -0.049 | 0.020 | 0.022 |
| 92 | Right | Default | Temporal | 0.207 | 0.242 | 0.189 | 0.187 | -0.385 | 0.016 | 0.000 |
| 93 | Right | Default | Temporal | 0.232 | 0.263 | 0.208 | 0.202 | -0.346 | 0.016 | 0.000 |
| 95 | Right | Default | Ventral Prefrontal Cortex | 0.081 | 0.103 | 0.079 | 0.094 | -0.081 | 0.019 | 0.000 |
| 97 | Right | Default | Dorsomedial Prefrontal Cortex | 0.093 | 0.118 | 0.066 | 0.066 | -0.302 | 0.019 | 0.000 |
| 98 | Right | Default | Dorsomedial Prefrontal Cortex | 0.086 | 0.103 | 0.062 | 0.069 | -0.119 | 0.019 | 0.000 |
| 100 | Left | Default | Precuneus/Posterior Cingulate Cortex | 0.100 | 0.110 | 0.091 | 0.096 | -0.061 | 0.019 | 0.002 |

Subcortical Results

| Parcel No. | Hemisphere | Region | $ISC_{hh.s}$ | $ISC_{ll.s}$ | $ISC_{hh.ns}$ | $ISC_{ll.ns}$ | β | *SE* | *p* |
|---|---|---|---|---|---|---|---|---|---|
| 101 | Left | Thalamus | 0.024 | 0.023 | 0.023 | 0.024 | 0.047 | 0.022 | 0.044 |
| 102 | Left | Caudate | 0.027 | 0.028 | 0.024 | 0.029 | 0.076 | 0.021 | 0.001 |
| 104 | Left | Pallidum | 0.004 | 0.005 | 0.002 | 0.005 | 0.065 | 0.023 | 0.006 |
| 106 | Left | Amygdala | 0.043 | 0.050 | 0.041 | 0.044 | -0.080 | 0.021 | 0.000 |
| 107 | Left | Accumbens | 0.009 | 0.008 | 0.008 | 0.011 | 0.077 | 0.022 | 0.001 |
| 108 | Left | Thalamus | 0.034 | 0.036 | 0.029 | 0.034 | 0.056 | 0.021 | 0.013 |
| 110 | Right | Putamen | 0.018 | 0.023 | 0.018 | 0.019 | -0.071 | 0.022 | 0.002 |
| 113 | Right | Amygdala | 0.054 | 0.062 | 0.050 | 0.048 | -0.178 | 0.021 | 0.000 |
| 114 | Right | Accumbens | 0.011 | 0.008 | 0.011 | 0.012 | 0.089 | 0.022 | 0.000 |

*Note.* This table corresponds to Fig. 3a in the manuscript. Parcel numbers for cortical regions correspond to the Schaefer 100 cortical parcellation (Schaefer et al., 2018), whereas parcel numbers for subcortical regions are arbitrary numbers assigned by researchers. Component names are from the labels provided by Yeo et al., (2011). $ISC_{hh.s}$ and $ISC_{ll.s}$ indicates mean ISC of the high-loneliness pairs or the low-loneliness pairs at the social context, respectively. $ISC_{hh.ns}$ and $ISC_{ll.ns}$ signifies mean ISC of the high-loneliness pairs or the low-loneliness pairs at the nonsocial context, respectively. Although linear mixed effects modelling considers the fully crossed random effects in comparing differences in the ISCs between groups for each context, we provide mean ISCs of each group in each context to enhance interpretability and facilitate direct comparison. **β** indicates standardized coefficients analyzed with the contrast testing whether ISC difference between high-loneliness pairs and low-loneliness pairs ($ISC_{hh} > ISC_{ll}$) is significantly different in social compared to nonsocial contexts. *SE* signifies standard error, and *p*-values were corrected for multiple comparisons using a false discovery rate at $p < .05$. Vis = Visual, SomMot = Somatomotor, DorsAttn = Dorsal Attention, SalVentAttn = Salience/Ventral Attention, Cont = Control.

**Table S6.** Brain regions showing significant interaction effects between loneliness and context ($ISC_{hh}$ > $ISC_{hl}$).

| Cortical Results | | | | | | | | | | |
|---|---|---|---|---|---|---|---|---|---|---|
| Parcel No. | Hemisphere | Network | Component | $ISC_{hh.s}$ | $ISC_{hl.s}$ | $ISC_{hh.ns}$ | $ISC_{hl.ns}$ | β | *SE* | *p* |
| 1 | Left | Vis | Visual | 0.224 | 0.229 | 0.165 | 0.173 | 0.028 | 0.013 | 0.045 |
| 2 | Left | Vis | Visual | 0.291 | 0.299 | 0.261 | 0.264 | -0.046 | 0.012 | 0.000 |
| 3 | Left | Vis | Visual | 0.308 | 0.310 | 0.221 | 0.229 | 0.048 | 0.011 | 0.000 |
| 4 | Left | Vis | Visual | 0.206 | 0.218 | 0.265 | 0.268 | -0.063 | 0.013 | 0.000 |
| 5 | Left | Vis | Visual | 0.291 | 0.304 | 0.317 | 0.324 | -0.043 | 0.011 | 0.001 |
| 6 | Left | Vis | Visual | 0.234 | 0.240 | 0.179 | 0.182 | -0.030 | 0.012 | 0.029 |
| 7 | Left | Vis | Visual | 0.318 | 0.327 | 0.294 | 0.298 | -0.038 | 0.011 | 0.002 |
| 8 | Left | Vis | Visual | 0.361 | 0.364 | 0.246 | 0.253 | 0.030 | 0.011 | 0.014 |
| 9 | Left | Vis | Visual | 0.212 | 0.208 | 0.166 | 0.173 | 0.090 | 0.014 | 0.000 |
| 10 | Left | SomMot | Somatomotor | 0.353 | 0.364 | 0.320 | 0.315 | -0.137 | 0.011 | 0.000 |
| 11 | Left | SomMot | Somatomotor | 0.054 | 0.054 | 0.057 | 0.053 | -0.059 | 0.017 | 0.001 |
| 13 | Left | SomMot | Somatomotor | 0.015 | 0.016 | 0.018 | 0.014 | -0.057 | 0.019 | 0.008 |
| 14 | Left | SomMot | Somatomotor | 0.021 | 0.021 | 0.027 | 0.023 | -0.051 | 0.019 | 0.018 |
| 15 | Left | SomMot | Somatomotor | 0.030 | 0.030 | 0.036 | 0.030 | -0.084 | 0.019 | 0.000 |
| 17 | Left | DorsAttn | Posterior | 0.151 | 0.147 | 0.157 | 0.160 | 0.070 | 0.014 | 0.000 |
| 18 | Left | DorsAttn | Posterior | 0.265 | 0.278 | 0.169 | 0.176 | -0.046 | 0.012 | 0.000 |
| 21 | Left | DorsAttn | Posterior | 0.168 | 0.174 | 0.162 | 0.162 | -0.048 | 0.013 | 0.001 |
| 26 | Left | SalVentAttn | Frontal Operculum/Insula | 0.028 | 0.026 | 0.034 | 0.038 | 0.082 | 0.019 | 0.000 |
| 27 | Left | SalVentAttn | Lateral Prefrontal Cortex | 0.050 | 0.046 | 0.040 | 0.044 | 0.101 | 0.019 | 0.000 |
| 28 | Left | SalVentAttn | Medial | 0.023 | 0.023 | 0.026 | 0.031 | 0.075 | 0.019 | 0.000 |
| 29 | Left | SalVentAttn | Medial | 0.116 | 0.111 | 0.089 | 0.089 | 0.058 | 0.016 | 0.001 |
| 32 | Left | Limbic | Temporal Pole | 0.085 | 0.089 | 0.078 | 0.079 | -0.037 | 0.017 | 0.048 |
| 33 | Left | Limbic | Temporal Pole | 0.078 | 0.085 | 0.044 | 0.046 | -0.072 | 0.017 | 0.000 |
| 35 | Left | Cont | Lateral Prefrontal Cortex | 0.063 | 0.063 | 0.043 | 0.049 | 0.079 | 0.018 | 0.000 |
| 36 | Left | Cont | Precuneus | 0.101 | 0.100 | 0.094 | 0.101 | 0.103 | 0.017 | 0.000 |
| 37 | Left | Cont | Cingulate Cortex | 0.067 | 0.067 | 0.059 | 0.064 | 0.064 | 0.018 | 0.001 |
| 38 | Left | Default | Temporal | 0.215 | 0.228 | 0.162 | 0.165 | -0.099 | 0.014 | 0.000 |
| 39 | Left | Default | Temporal | 0.240 | 0.259 | 0.175 | 0.177 | -0.151 | 0.013 | 0.000 |
| 40 | Left | Default | Parietal | 0.188 | 0.198 | 0.173 | 0.177 | -0.063 | 0.016 | 0.000 |
| 43 | Left | Default | Prefrontal Cortex | 0.112 | 0.114 | 0.050 | 0.056 | 0.043 | 0.016 | 0.015 |
| 46 | Left | Default | Prefrontal Cortex | 0.091 | 0.101 | 0.046 | 0.050 | -0.079 | 0.017 | 0.000 |

| | | | | | | | | | | |
|---|---|---|---|---|---|---|---|---|---|---|
| 50 | Left | Default | Precuneus/Posterior Cingulate Cortex | 0.098 | 0.106 | 0.079 | 0.084 | -0.041 | 0.017 | 0.029 |
| 54 | Right | Vis | Visual | 0.220 | 0.231 | 0.273 | 0.278 | -0.043 | 0.012 | 0.001 |
| 55 | Right | Vis | Visual | 0.261 | 0.267 | 0.214 | 0.225 | 0.032 | 0.013 | 0.022 |
| 56 | Right | Vis | Visual | 0.231 | 0.228 | 0.183 | 0.187 | 0.053 | 0.013 | 0.000 |
| 58 | Right | Vis | Visual | 0.275 | 0.274 | 0.198 | 0.209 | 0.091 | 0.012 | 0.000 |
| 59 | Right | SomMot | Somatomotor | 0.309 | 0.317 | 0.285 | 0.280 | -0.109 | 0.012 | 0.000 |
| 60 | Right | SomMot | Somatomotor | 0.027 | 0.025 | 0.029 | 0.024 | -0.058 | 0.019 | 0.006 |
| 62 | Right | SomMot | Somatomotor | 0.013 | 0.013 | 0.016 | 0.012 | -0.060 | 0.020 | 0.005 |
| 64 | Right | SomMot | Somatomotor | 0.021 | 0.021 | 0.030 | 0.023 | -0.083 | 0.019 | 0.000 |
| 65 | Right | SomMot | Somatomotor | 0.088 | 0.091 | 0.065 | 0.063 | -0.055 | 0.015 | 0.001 |
| 66 | Right | SomMot | Somatomotor | 0.033 | 0.032 | 0.041 | 0.033 | -0.093 | 0.019 | 0.000 |
| 70 | Right | DorsAttn | Posterior | 0.268 | 0.280 | 0.152 | 0.161 | -0.030 | 0.012 | 0.023 |
| 71 | Right | DorsAttn | Posterior | 0.180 | 0.190 | 0.130 | 0.131 | -0.079 | 0.013 | 0.000 |
| 74 | Right | SalVentAttn | Temporal Occipital Parietal | 0.242 | 0.253 | 0.205 | 0.202 | -0.123 | 0.013 | 0.000 |
| 75 | Right | SalVentAttn | Temporal Occipital Parietal | 0.123 | 0.119 | 0.132 | 0.133 | 0.056 | 0.014 | 0.000 |
| 76 | Right | SalVentAttn | Frontal Operculum/Insula | 0.036 | 0.033 | 0.041 | 0.044 | 0.085 | 0.019 | 0.000 |
| 78 | Right | SalVentAttn | Medial | 0.028 | 0.027 | 0.027 | 0.030 | 0.046 | 0.019 | 0.029 |
| 79 | Right | Limbic | Orbitofrontal Cortex | 0.021 | 0.021 | 0.011 | 0.014 | 0.071 | 0.019 | 0.001 |
| 80 | Right | Limbic | Temporal Pole | 0.090 | 0.096 | 0.076 | 0.077 | -0.073 | 0.017 | 0.000 |
| 81 | Right | Cont | Parietal | 0.091 | 0.093 | 0.102 | 0.110 | 0.073 | 0.016 | 0.000 |
| 82 | Right | Cont | Parietal | 0.087 | 0.095 | 0.079 | 0.083 | -0.050 | 0.017 | 0.008 |
| 83 | Right | Cont | Lateral Prefrontal Cortex | 0.053 | 0.059 | 0.059 | 0.062 | -0.046 | 0.018 | 0.022 |
| 85 | Right | Cont | Lateral Prefrontal Cortex | 0.057 | 0.056 | 0.057 | 0.062 | 0.065 | 0.018 | 0.001 |
| 86 | Right | Cont | Lateral Prefrontal Cortex | 0.051 | 0.062 | 0.069 | 0.074 | -0.077 | 0.018 | 0.000 |
| 87 | Right | Cont | Cingulate Cortex | 0.066 | 0.065 | 0.067 | 0.070 | 0.057 | 0.018 | 0.004 |
| 88 | Right | Cont | Medial Posterior Prefrontal Cortex | 0.027 | 0.028 | 0.035 | 0.043 | 0.087 | 0.019 | 0.000 |
| 92 | Right | Default | Temporal | 0.207 | 0.223 | 0.189 | 0.188 | -0.177 | 0.014 | 0.000 |
| 93 | Right | Default | Temporal | 0.232 | 0.247 | 0.208 | 0.206 | -0.155 | 0.014 | 0.000 |
| 94 | Right | Default | Ventral Prefrontal Cortex | 0.028 | 0.030 | 0.035 | 0.040 | 0.040 | 0.019 | 0.048 |
| 97 | Right | Default | Dorsomedial Prefrontal Cortex | 0.093 | 0.106 | 0.066 | 0.068 | -0.124 | 0.017 | 0.000 |
| 98 | Right | Default | Dorsomedial Prefrontal Cortex | 0.086 | 0.095 | 0.062 | 0.066 | -0.059 | 0.017 | 0.001 |

Subcortical Results

| Parcel No. | Hemisphere | Region | $ISC_{hh.s}$ | $ISC_{hl.s}$ | $ISC_{hh.ns}$ | $ISC_{hl.ns}$ | $\beta$ | *SE* | *p* |
|---|---|---|---|---|---|---|---|---|---|
| 102 | Left | Caudate | 0.027 | 0.028 | 0.024 | 0.028 | 0.058 | 0.019 | 0.005 |
| 107 | Left | Accumbens | 0.009 | 0.009 | 0.008 | 0.010 | 0.046 | 0.020 | 0.032 |

| | | | | | | | | | |
|---|---|---|---|---|---|---|---|---|---|
| 108 | Left | Thalamus | 0.034 | 0.035 | 0.029 | 0.032 | 0.048 | 0.019 | 0.022 |
| 112 | Right | Hippocampus | 0.042 | 0.042 | 0.040 | 0.043 | 0.045 | 0.019 | 0.029 |
| 113 | Right | Amygdala | 0.054 | 0.057 | 0.050 | 0.050 | -0.058 | 0.018 | 0.004 |
| 114 | Right | Accumbens | 0.011 | 0.009 | 0.011 | 0.012 | 0.069 | 0.020 | 0.001 |

*Note.* This table corresponds to Fig. 3b in the manuscript. Parcel numbers for cortical regions correspond to the Schaefer 100 cortical parcellation (Schaefer et al., 2018), whereas parcel numbers for subcortical regions are arbitrary numbers assigned by researchers. Component names are from the labels provided by Yeo et al., (2011). $ISC_{hh.s}$ and $ISC_{hl.s}$ indicates mean ISC of the high-loneliness pairs or the mismatched pairs at the social context, respectively. $ISC_{hh.ns}$ and $ISC_{hl.ns}$ signifies mean ISC of the high-loneliness pairs or the mismatched pairs at the nonsocial context, respectively. Although linear mixed effects modelling considers the fully crossed random effects in comparing differences in the ISCs between groups for each context, we provide mean ISCs of each group in each context to enhance interpretability and facilitate direct comparison. **β** indicates standardized coefficients analyzed with the contrast testing whether ISC difference between high-loneliness pairs and mismatched pairs ($ISC_{hh} > ISC_{hl}$) is significantly different in social compared to nonsocial contexts. *SE* signifies standard error, and *p*-values were corrected for multiple comparisons using a false discovery rate at $p < .05$. Vis = Visual, SomMot = Somatomotor, DorsAttn = Dorsal Attention, SalVentAttn = Salience/Ventral Attention, Cont = Control.

**Table S7.** Brain regions showing significant interaction effects between loneliness and context ($ISC_{hl}$ > $ISC_{ll}$).

| | | | Cortical Results | | | | | | | |
|---|---|---|---|---|---|---|---|---|---|---|
| Parcel No. | Hemisphere | Network | Component | $ISC_{hl.s}$ | $ISC_{ll.s}$ | $ISC_{hl.ns}$ | $ISC_{ll.ns}$ | β | *SE* | *p* |
| 2 | Left | Vis | Visual | 0.299 | 0.310 | 0.264 | 0.268 | -0.052 | 0.012 | 0.000 |
| 3 | Left | Vis | Visual | 0.310 | 0.315 | 0.229 | 0.237 | 0.026 | 0.011 | 0.028 |
| 4 | Left | Vis | Visual | 0.218 | 0.231 | 0.268 | 0.274 | -0.058 | 0.012 | 0.000 |
| 5 | Left | Vis | Visual | 0.304 | 0.319 | 0.324 | 0.332 | -0.054 | 0.011 | 0.000 |
| 6 | Left | Vis | Visual | 0.240 | 0.248 | 0.182 | 0.183 | -0.059 | 0.012 | 0.000 |
| 7 | Left | Vis | Visual | 0.327 | 0.337 | 0.298 | 0.303 | -0.043 | 0.011 | 0.000 |
| 8 | Left | Vis | Visual | 0.364 | 0.369 | 0.253 | 0.262 | 0.032 | 0.011 | 0.008 |
| 9 | Left | Vis | Visual | 0.208 | 0.208 | 0.173 | 0.178 | 0.051 | 0.013 | 0.000 |
| 10 | Left | SomMot | Somatomotor | 0.364 | 0.375 | 0.315 | 0.311 | -0.121 | 0.011 | 0.000 |
| 11 | Left | SomMot | Somatomotor | 0.054 | 0.056 | 0.053 | 0.048 | -0.112 | 0.017 | 0.000 |
| 12 | Left | SomMot | Somatomotor | 0.049 | 0.050 | 0.042 | 0.038 | -0.076 | 0.018 | 0.000 |
| 13 | Left | SomMot | Somatomotor | 0.016 | 0.016 | 0.014 | 0.008 | -0.087 | 0.019 | 0.000 |
| 14 | Left | SomMot | Somatomotor | 0.021 | 0.022 | 0.023 | 0.018 | -0.068 | 0.019 | 0.001 |
| 15 | Left | SomMot | Somatomotor | 0.030 | 0.031 | 0.030 | 0.026 | -0.070 | 0.018 | 0.000 |
| 17 | Left | DorsAttn | Posterior | 0.147 | 0.147 | 0.160 | 0.163 | 0.032 | 0.013 | 0.032 |
| 18 | Left | DorsAttn | Posterior | 0.278 | 0.290 | 0.176 | 0.183 | -0.051 | 0.012 | 0.000 |
| 21 | Left | DorsAttn | Posterior | 0.174 | 0.181 | 0.162 | 0.163 | -0.048 | 0.013 | 0.000 |
| 24 | Left | SalVentAttn | Parietal Operculum | 0.086 | 0.090 | 0.116 | 0.115 | -0.061 | 0.016 | 0.000 |
| 31 | Left | Limbic | Orbitofrontal Cortex | 0.025 | 0.027 | 0.016 | 0.020 | 0.065 | 0.018 | 0.001 |
| 32 | Left | Limbic | Temporal Pole | 0.089 | 0.096 | 0.079 | 0.080 | -0.087 | 0.016 | 0.000 |
| 35 | Left | Cont | Lateral Prefrontal Cortex | 0.063 | 0.063 | 0.049 | 0.055 | 0.073 | 0.018 | 0.000 |
| 36 | Left | Cont | Precuneus | 0.100 | 0.100 | 0.101 | 0.108 | 0.081 | 0.017 | 0.000 |
| 38 | Left | Default | Temporal | 0.228 | 0.245 | 0.165 | 0.167 | -0.137 | 0.014 | 0.000 |
| 39 | Left | Default | Temporal | 0.259 | 0.279 | 0.177 | 0.177 | -0.179 | 0.013 | 0.000 |
| 40 | Left | Default | Parietal | 0.198 | 0.211 | 0.177 | 0.180 | -0.091 | 0.015 | 0.000 |
| 42 | Left | Default | Prefrontal Cortex | 0.058 | 0.063 | 0.043 | 0.045 | -0.043 | 0.017 | 0.026 |
| 43 | Left | Default | Prefrontal Cortex | 0.114 | 0.114 | 0.056 | 0.060 | 0.042 | 0.016 | 0.015 |
| 44 | Left | Default | Prefrontal Cortex | 0.035 | 0.038 | 0.036 | 0.035 | -0.047 | 0.018 | 0.021 |
| 46 | Left | Default | Prefrontal Cortex | 0.101 | 0.110 | 0.050 | 0.051 | -0.092 | 0.016 | 0.000 |
| 50 | Left | Default | Precuneus/Posterior Cingulate Cortex | 0.106 | 0.115 | 0.084 | 0.089 | -0.045 | 0.016 | 0.012 |
| 54 | Right | Vis | Visual | 0.231 | 0.242 | 0.278 | 0.285 | -0.030 | 0.012 | 0.023 |

| | | | | | | | | | | |
|---|---|---|---|---|---|---|---|---|---|---|
| 56 | Right | Vis | Visual | 0.228 | 0.226 | 0.187 | 0.189 | 0.044 | 0.012 | 0.001 |
| 58 | Right | Vis | Visual | 0.274 | 0.275 | 0.209 | 0.218 | 0.067 | 0.012 | 0.000 |
| 59 | Right | SomMot | Somatomotor | 0.317 | 0.325 | 0.280 | 0.276 | -0.095 | 0.011 | 0.000 |
| 60 | Right | SomMot | Somatomotor | 0.025 | 0.026 | 0.024 | 0.020 | -0.078 | 0.019 | 0.000 |
| 61 | Right | SomMot | Somatomotor | 0.051 | 0.051 | 0.033 | 0.030 | -0.051 | 0.018 | 0.011 |
| 62 | Right | SomMot | Somatomotor | 0.013 | 0.013 | 0.012 | 0.007 | -0.067 | 0.019 | 0.001 |
| 63 | Right | SomMot | Somatomotor | 0.022 | 0.023 | 0.022 | 0.019 | -0.053 | 0.019 | 0.011 |
| 65 | Right | SomMot | Somatomotor | 0.091 | 0.093 | 0.063 | 0.062 | -0.035 | 0.015 | 0.040 |
| 66 | Right | SomMot | Somatomotor | 0.032 | 0.033 | 0.033 | 0.030 | -0.051 | 0.018 | 0.012 |
| 67 | Right | DorsAttn | Posterior | 0.265 | 0.282 | 0.200 | 0.210 | -0.059 | 0.013 | 0.000 |
| 70 | Right | DorsAttn | Posterior | 0.280 | 0.291 | 0.161 | 0.168 | -0.037 | 0.012 | 0.003 |
| 71 | Right | DorsAttn | Posterior | 0.190 | 0.199 | 0.131 | 0.132 | -0.062 | 0.013 | 0.000 |
| 74 | Right | SalVentAttn | Temporal Occipital Parietal | 0.253 | 0.265 | 0.202 | 0.199 | -0.145 | 0.013 | 0.000 |
| 77 | Right | SalVentAttn | Medial | 0.101 | 0.105 | 0.076 | 0.076 | -0.041 | 0.015 | 0.016 |
| 78 | Right | SalVentAttn | Medial | 0.027 | 0.031 | 0.030 | 0.031 | -0.048 | 0.018 | 0.020 |
| 79 | Right | Limbic | Orbitofrontal Cortex | 0.021 | 0.019 | 0.014 | 0.017 | 0.089 | 0.019 | 0.000 |
| 80 | Right | Limbic | Temporal Pole | 0.096 | 0.106 | 0.077 | 0.076 | -0.164 | 0.016 | 0.000 |
| 82 | Right | Cont | Parietal | 0.095 | 0.105 | 0.083 | 0.086 | -0.071 | 0.017 | 0.000 |
| 83 | Right | Cont | Lateral Prefrontal Cortex | 0.059 | 0.066 | 0.062 | 0.059 | -0.115 | 0.018 | 0.000 |
| 84 | Right | Cont | Lateral Prefrontal Cortex | 0.078 | 0.085 | 0.065 | 0.065 | -0.074 | 0.017 | 0.000 |
| 86 | Right | Cont | Lateral Prefrontal Cortex | 0.062 | 0.074 | 0.074 | 0.076 | -0.118 | 0.017 | 0.000 |
| 90 | Right | Default | Parietal | 0.089 | 0.098 | 0.103 | 0.105 | -0.077 | 0.016 | 0.000 |
| 92 | Right | Default | Temporal | 0.223 | 0.242 | 0.188 | 0.187 | -0.208 | 0.014 | 0.000 |
| 93 | Right | Default | Temporal | 0.247 | 0.263 | 0.206 | 0.202 | -0.191 | 0.013 | 0.000 |
| 95 | Right | Default | Ventral Prefrontal Cortex | 0.092 | 0.103 | 0.089 | 0.094 | -0.071 | 0.016 | 0.000 |
| 97 | Right | Default | Dorsomedial Prefrontal Cortex | 0.106 | 0.118 | 0.068 | 0.066 | -0.178 | 0.016 | 0.000 |
| 98 | Right | Default | Dorsomedial Prefrontal Cortex | 0.095 | 0.103 | 0.066 | 0.069 | -0.060 | 0.016 | 0.001 |

Subcortical Results

| Parcel No. | Hemisphere | Region | $ISC_{hl.s}$ | $ISC_{ll.s}$ | $ISC_{hl.ns}$ | $ISC_{ll.ns}$ | β | *SE* | *p* |
|---|---|---|---|---|---|---|---|---|---|
| 105 | Left | Hippocampus | 0.038 | 0.040 | 0.037 | 0.037 | -0.041 | 0.018 | 0.047 |
| 106 | Left | Amygdala | 0.046 | 0.050 | 0.043 | 0.044 | -0.059 | 0.018 | 0.002 |
| 110 | Right | Putamen | 0.020 | 0.023 | 0.019 | 0.019 | -0.059 | 0.019 | 0.004 |
| 113 | Right | Amygdala | 0.057 | 0.062 | 0.050 | 0.048 | -0.120 | 0.018 | 0.000 |

*Note.* This table corresponds to Fig. 3c in the manuscript. Parcel numbers for cortical regions correspond to the Schaefer 100 cortical parcellation (Schaefer et al., 2018), whereas parcel numbers for subcortical regions are arbitrary numbers assigned by researchers. Component names are from the labels provided by Yeo et al., (2011). $ISC_{hl.s}$

and $ISC_{ll.s}$ indicates mean ISC of the mismatched pairs or the low-loneliness pairs at the social context, respectively. $ISC_{hl.ns}$ and $ISC_{ll.ns}$ signifies mean ISC of the mismatched pairs or the low-loneliness pairs at the nonsocial context, respectively. Although linear mixed effects modelling considers the fully crossed random effects in comparing differences in the ISCs between groups for each context, we provide mean ISCs of each group in each context to enhance interpretability and facilitate direct comparison. **β** indicates standardized coefficients analyzed with the contrast testing whether ISC difference between mismatched pairs and low-loneliness pairs ($ISC_{hl} > ISC_{ll}$) is significantly different in social compared to nonsocial contexts. *SE* signifies standard error, and *p*-values were corrected for multiple comparisons using a false discovery rate at $p < .05$. Vis = Visual, SomMot = Somatomotor, DorsAttn = Dorsal Attention, SalVentAttn = Salience/Ventral Attention, Cont = Control.

**Table S8.** Brain regions showing significant effect of loneliness in the social context.

| | | Contrast: $ISC_{hh} > ISC_{ll}$ | | | | | |
|---|---|---|---|---|---|---|---|
| Parcel No. | Hemisphere | Region | $ISC_{hh}$ | $ISC_{ll}$ | β | *SE* | *p* |
| 38 | Left | MTG | 0.215 | 0.245 | -0.279 | 0.095 | 0.048 |
| 39 | Left | MTG | 0.240 | 0.279 | -0.342 | 0.096 | 0.011 |
| 46 | Left | dlPFC | 0.091 | 0.110 | -0.217 | 0.072 | 0.048 |
| 80 | Right | TmpPole | 0.090 | 0.106 | -0.232 | 0.079 | 0.048 |
| 86 | Right | FEF | 0.051 | 0.074 | -0.280 | 0.065 | 0.002 |
| 92 | Right | TmpPole | 0.207 | 0.242 | -0.362 | 0.101 | 0.011 |
| 95 | Right | BrTr | 0.081 | 0.103 | -0.253 | 0.080 | 0.034 |
| 97 | Right | dmPFC | 0.093 | 0.118 | -0.297 | 0.074 | 0.003 |
| | | Contrast: $ISC_{hh} > ISC_{hl}$ | | | | | |
| Parcel No. | Hemisphere | Region | $ISC_{hh}$ | $ISC_{hl}$ | β | *SE* | *p* |
| 39 | Left | MTG | 0.240 | 0.259 | -0.167 | 0.049 | 0.023 |
| 46 | Left | dlPFC | 0.091 | 0.101 | -0.113 | 0.037 | 0.048 |
| 86 | Right | FEF | 0.051 | 0.062 | -0.136 | 0.034 | 0.005 |
| 92 | Right | TemPole | 0.207 | 0.223 | -0.163 | 0.051 | 0.043 |
| 95 | Right | BrTr | 0.081 | 0.092 | -0.126 | 0.041 | 0.047 |
| 97 | Right | dmPFC | 0.093 | 0.106 | -0.151 | 0.038 | 0.005 |
| | | Contrast: $ISC_{hl} > ISC_{ll}$ | | | | | |
| Parcel No. | Hemisphere | Region | $ISC_{hl}$ | $ISC_{ll}$ | β | *SE* | *p* |
| 38 | Left | MTG | 0.228 | 0.245 | -0.157 | 0.048 | 0.022 |
| 39 | Left | MTG | 0.259 | 0.279 | -0.175 | 0.049 | 0.008 |
| 46 | Left | dlPFC | 0.101 | 0.110 | -0.104 | 0.037 | 0.047 |
| 67 | Right | VAC | 0.265 | 0.282 | -0.149 | 0.053 | 0.047 |
| 80 | Right | TmpPole | 0.096 | 0.106 | -0.151 | 0.040 | 0.005 |
| 82 | Right | AngGyr | 0.095 | 0.105 | -0.107 | 0.037 | 0.047 |
| 83 | Right | aPFC | 0.059 | 0.066 | -0.088 | 0.032 | 0.047 |
| 86 | Right | FEF | 0.062 | 0.074 | -0.144 | 0.034 | 0.003 |
| 92 | Right | TmpPole | 0.223 | 0.242 | -0.199 | 0.051 | 0.005 |
| 93 | Right | STG | 0.247 | 0.263 | -0.153 | 0.053 | 0.047 |
| 95 | Right | BrTr | 0.092 | 0.103 | -0.127 | 0.041 | 0.032 |
| 97 | Right | dmPFC | 0.106 | 0.118 | -0.146 | 0.038 | 0.005 |
| 113 | Right | Amg | 0.057 | 0.062 | -0.091 | 0.032 | 0.047 |

*Note.* This table corresponds to Fig. 4 in the manuscript. Parcel numbers for cortical regions correspond to the Schaefer 100 cortical parcellation (Schaefer et al., 2018), whereas parcel numbers for subcortical regions are arbitrary numbers assigned by researchers. Although linear mixed effects modelling considers the fully crossed random effects in comparing differences in the ISCs between groups, we provide mean ISCs of each group to enhance interpretability and facilitate direct comparison. **β** indicates standardized coefficients analyzed with each contrast testing whether ISCs between groups are significantly different in the social context. *SE* signifies standard error, and *p*-values were corrected for multiple comparisons using a false discovery rate at $p < .05$. MTG = Medial temporal gyrus, dlPFC = dorsolateral prefrontal cortex, TmpPole = Temporal pole, FEF = Frontal eye fields, BrTr = Broca's Triangle, dmPFC = dorsomedial prefrontal cortex, VAC = Visual association cortex, AngGyr = Angular gyrus, aPFC = anterior prefrontal cortex, STG = Superior temporal gyrus, Amg = Amygdala.

**Table S9.** Brain regions showing significant effect of loneliness in the nonsocial context.

| | | Contrast: $ISC_{hh} > ISC_{ll}$ | | | | | |
|---|---|---|---|---|---|---|---|
| Parcel No. | Hemisphere | Region | $ISC_{hh}$ | $ISC_{ll}$ | β | *SE* | *p* |
| 13 | Left | PreMot + SuppMot | 0.018 | 0.008 | 0.132 | 0.032 | 0.001 |
| 14 | Left | PrimMot | 0.027 | 0.018 | 0.117 | 0.037 | 0.023 |
| 15 | Left | PrimMot | 0.036 | 0.026 | 0.145 | 0.045 | 0.019 |
| 60 | Right | Ins | 0.029 | 0.020 | 0.149 | 0.040 | 0.004 |
| 62 | Right | PrimMot | 0.016 | 0.007 | 0.126 | 0.030 | 0.001 |
| 64 | Right | PrimMot | 0.030 | 0.019 | 0.156 | 0.037 | 0.001 |
| 66 | Right | PrimMot | 0.041 | 0.030 | 0.154 | 0.045 | 0.010 |
| 79 | Right | OFC | 0.011 | 0.017 | -0.116 | 0.039 | 0.036 |
| 88 | Right | mpPFC | 0.035 | 0.045 | -0.139 | 0.047 | 0.036 |
| 104 | Left | Pal | 0.002 | 0.005 | -0.088 | 0.022 | 0.001 |
| | | Contrast: $ISC_{hh} > ISC_{hl}$ | | | | | |
| Parcel No. | Hemisphere | Region | $ISC_{hh}$ | $ISC_{hl}$ | β | *SE* | *p* |
| 15 | Left | PrimMot | 0.036 | 0.030 | 0.092 | 0.025 | 0.005 |
| 28 | Left | dACC | 0.026 | 0.031 | -0.066 | 0.022 | 0.037 |
| 60 | Right | Ins | 0.029 | 0.024 | 0.084 | 0.023 | 0.005 |
| 62 | Right | PrimMot | 0.016 | 0.012 | 0.056 | 0.019 | 0.037 |
| 64 | Right | PrimMot | 0.030 | 0.023 | 0.095 | 0.021 | 0.001 |
| 66 | Right | PrimMot | 0.041 | 0.033 | 0.108 | 0.025 | 0.001 |
| 88 | Right | mpPFC | 0.035 | 0.043 | -0.108 | 0.026 | 0.001 |
| 95 | Right | BrTr | 0.079 | 0.089 | -0.117 | 0.041 | 0.050 |
| 102 | Left | Caud | 0.024 | 0.028 | -0.076 | 0.027 | 0.050 |
| 104 | Left | Pal | 0.002 | 0.004 | -0.056 | 0.016 | 0.006 |
| 107 | Left | NAcc | 0.008 | 0.010 | -0.051 | 0.018 | 0.050 |
| | | Contrast: $ISC_{hl} > ISC_{ll}$ | | | | | |
| Parcel No. | Hemisphere | Region | $ISC_{hl}$ | $ISC_{ll}$ | β | *SE* | *p* |
| 13 | Left | PreMot + SuppMot | 0.014 | 0.008 | 0.081 | 0.019 | 0.003 |
| 31 | Left | OFC | 0.016 | 0.020 | -0.091 | 0.024 | 0.008 |
| 62 | Right | PrimMot | 0.012 | 0.007 | 0.069 | 0.018 | 0.008 |

*Note.* This table corresponds to Fig. 5 in the manuscript. Parcel numbers for cortical regions correspond to the Schaefer 100 cortical parcellation (Schaefer et al., 2018), whereas parcel numbers for subcortical regions are arbitrary numbers assigned by researchers. Although linear mixed effects modelling considers the fully crossed random effects in comparing differences in the ISCs between groups, we provide mean ISCs of each group to enhance interpretability and facilitate direct comparison. **β** indicates standardized coefficients analyzed with each contrast testing whether ISCs between groups are significantly different in the nonsocial context. *SE* signifies standard error, and *p*-values were corrected for multiple comparisons using a false discovery rate at $p < .05$. PreMot + SuppMot = Premotor and Supplementary cortex, PrimMot = Primary motor cortex, Ins = Insula, OFC = Orbitofrontal cortex, mpPFC = medial posterior prefrontal cortex, Pal = Pallidum, dACC = dorsal anterior cingulate cortex, BrTr = Broca's Triangle, Caud = Caudate, NAcc = Accumbens.

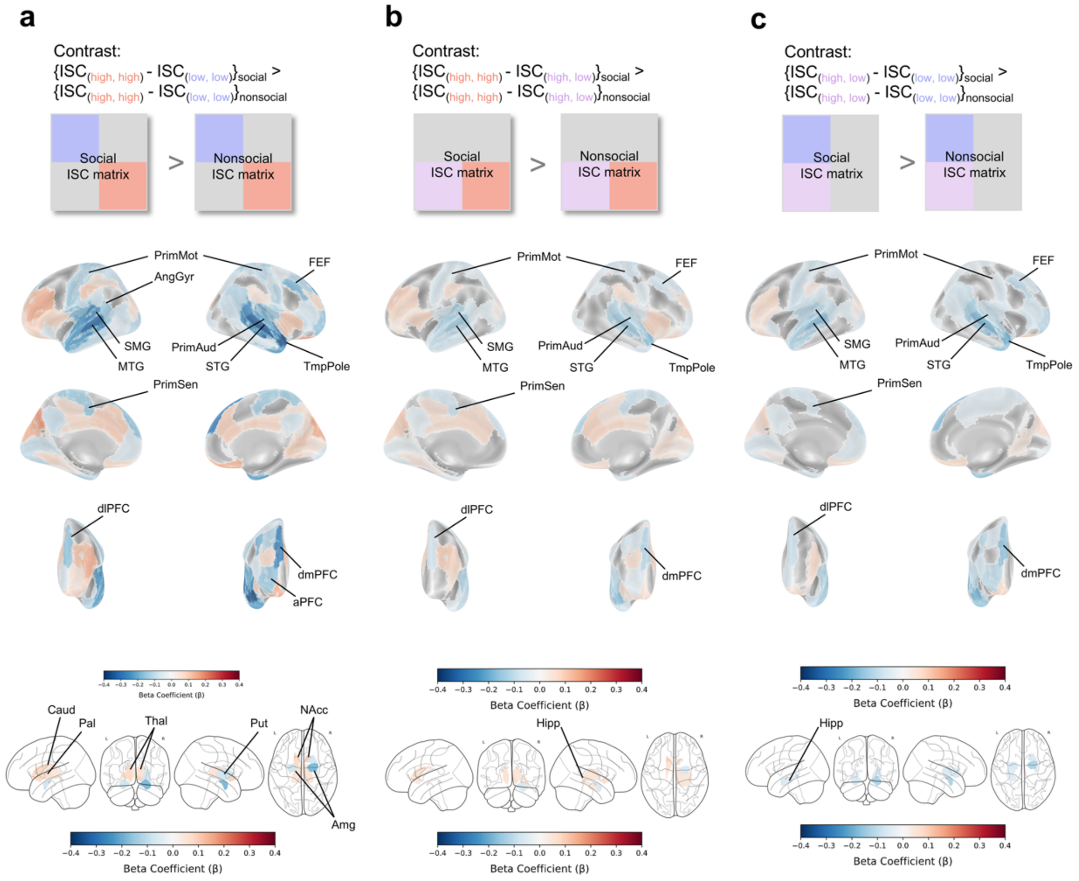


**Fig. S1.** Interaction effect between loneliness and context when controlling for covariates. **a.** Schematized matrices indicate a contrast testing whether ISC difference between high-loneliness pairs and low-loneliness pairs ($ISC_{hh} > ISC_{ll}$) is significantly different in social compared to nonsocial context (top panel). Cortical results (middle panel) and subcortical results (bottom panel) are shown with a false-discovery-rate (FDR) corrected significance threshold of $p < .05$. **b.** Schematized matrices signify a contrast testing whether ISC difference between high-loneliness pairs and mismatched pairs ($ISC_{hh} > ISC_{hl}$) is significantly different in social compared to nonsocial contexts (top panel). Cortical results (middle panel) and subcortical results (bottom panel) are shown with an FDR-corrected significance threshold of $p < .05$. **c.** Schematized matrices indicate a contrast testing whether ISC difference between mismatched pairs and low-loneliness pairs ($ISC_{hl} > ISC_{ll}$) is significantly different in social compared to nonsocial contexts (top panel). Cortical results (middle panel) and subcortical results (bottom panel) are shown with an FDR-corrected significance threshold of $p < .05$. Each social and nonsocial ISC matrix is simplified to aid identification of the contrast. Control variables used in the model were demographic similarities (age, gender, and race), kinship, friendship survey from the NIH Toolbox on emotion, and Achenbach adult self-report DSM-oriented depressive problems scale. For details of analysis with the model including covariates of no interest, see Methods section in the manuscript. PrimMot = Primary motor cortex, AngGyr = Angular gyrus, SMG = Supramarginal gyrus, MTG = Medial temporal gyrus, FEF = Frontal eye fields, PrimAud = Primary auditory cortex, STG = Superior temporal gyrus, TmpPole = Temporal pole, PrimSen = Primary sensory cortex, dlPFC = dorsolateral prefrontal cortex, dmPFC = dorsomedial prefrontal cortex, aPFC = anterior prefrontal cortex, Caud = Caudate, Pal = Pallidum, Thal = Thalamus, Put = Putamen, NAcc = Accumbens, Amg = Amygdala, Hipp = Hippocampus.

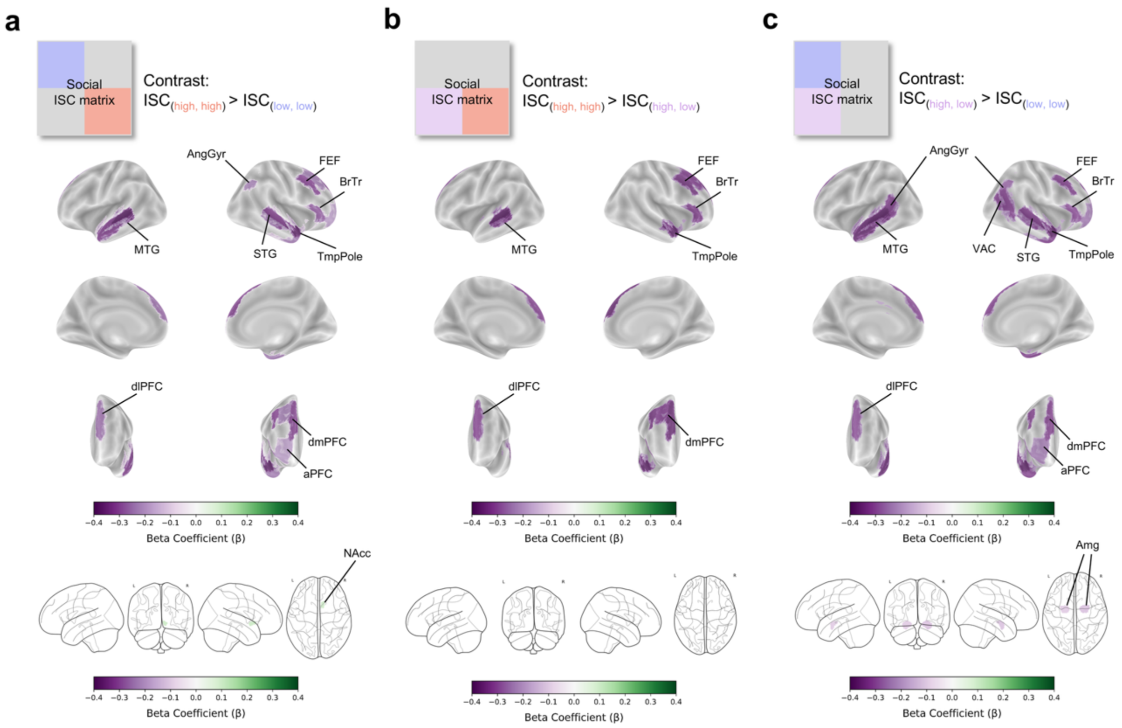


**Fig S2.** Simple main effect of loneliness in the social context while controlling for covariates. **a.** Schematized matrix indicates a contrast testing whether ISCs of high-loneliness pairs and low-loneliness pairs ($ISC_{hh} > ISC_{ll}$) are significantly different during viewing social context. (top panel). Cortical results (middle panel) and subcortical results (bottom panel) are shown with a false-discovery-rate (FDR) corrected significance threshold of $p < .05$. **b.** Schematized matrix signifies a contrast testing whether ISCs of high-loneliness pairs and mismatched pairs ($ISC_{hh} > ISC_{hl}$) are significantly different while viewing social scenes (top panel). Cortical results (middle panel) and subcortical results (bottom panel) are shown with an FDR-corrected significance threshold of $p < .05$. **c.** Schematized matrix indicates a contrast testing whether ISCs of mismatched pairs and low-loneliness pairs ($ISC_{hl} > ISC_{ll}$) are significantly different during viewing social context (top panel). Cortical results (middle panel) and subcortical results (bottom panel) are shown with an FDR-corrected significance threshold of $p < .05$. The social ISC matrix is simplified to aid identification of the contrast. Control variables used in the model were demographic similarities (age, gender, and race), kinship, friendship survey from the NIH Toolbox on emotion, and Achenbach adult self-report DSM-oriented depressive problems scale. For details of analysis with the model including covariates of no interest, see Methods section in the manuscript. MTG = Medial temporal gyrus, AngGyr = Angular gyrus, FEF = Frontal eye fields, BrTr = Broca's triangle, STG = Superior temporal gyrus, TmpPole = Temporal pole, dlPFC = dorsolateral prefrontal cortex, dmPFC = dorsomedial prefrontal cortex, aPFC = anterior prefrontal cortex, NAcc = Accumbens, VAC = Visual association cortex, Amg = Amygdala.

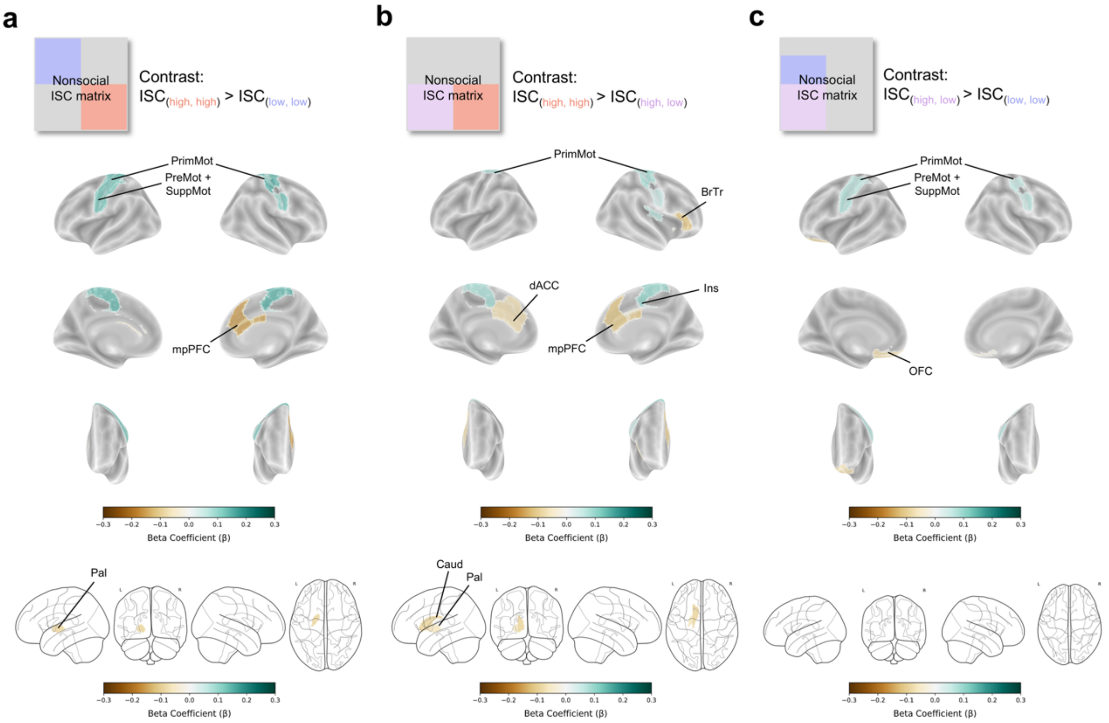


**Fig S3.** Simple main effect of loneliness in the nonsocial context while controlling for covariates. **a.** Schematized matrix indicates a contrast testing whether ISCs of high-loneliness pairs and low-loneliness pairs ($ISC_{hh} > ISC_{ll}$) are significantly different during viewing nonsocial context. (top panel). Cortical results (middle panel) and subcortical results (bottom panel) are shown with a false-discovery-rate (FDR) corrected significance threshold of $p < .05$. **b.** Schematized matrix signifies a contrast testing whether ISCs of high-loneliness pairs and mismatched pairs ($ISC_{hh} > ISC_{hl}$) are significantly different while viewing nonsocial scenes (top panel). Cortical results (middle panel) and subcortical results (bottom panel) are shown with an FDR-corrected significance threshold of $p < .05$. **c.** Schematized matrix indicates a contrast testing whether ISCs of mismatched pairs and low-loneliness pairs ($ISC_{hl} > ISC_{ll}$) are significantly different during viewing nonsocial context (top panel). Cortical results (middle panel) and subcortical results (bottom panel) are shown with an FDR-corrected significance threshold of $p < .05$. The nonsocial ISC matrix is simplified to aid identification of the contrast. Control variables used in the model were demographic similarities (age, gender, and race), kinship, friendship survey from the NIH Toolbox on emotion, and Achenbach adult self-report DSM-oriented depressive problems scale. For details of analysis with the model including covariates of no interest, see Methods section in the manuscript. PrimMot = Primary motor cortex, PreMot + SuppMot = Premotor and Supplementary motor cortex, mpPFC = medial posterior prefrontal cortex, Pal = Pallidum, BrTr = Broca's triangle, dACC = dorsal anterior cingulate cortex, Ins = Insula, Caud = Caudate, OFC = Orbitofrontal cortex.
